# Single-molecule measurements reveal that PARP1 condenses DNA by loop formation

**DOI:** 10.1101/2020.09.15.297887

**Authors:** Nicholas A. W. Bell, Philip J. Haynes, Katharina Brunner, Taiana Maia de Oliveira, Maria Flocco, Bart W. Hoogenboom, Justin E. Molloy

## Abstract

Poly(ADP-ribose) polymerase 1 (PARP1) is an abundant nuclear enzyme that plays important roles in DNA repair, chromatin organization and transcription regulation. Although binding and activation of PARP1 by DNA damage sites has been extensively studied, little is known about how PARP1 binds to long stretches of undamaged DNA and how it could shape chromatin architecture. Here, using a combination of single-molecule techniques including magnetic tweezers and atomic force microscopy, we show that PARP1 binds and condenses undamaged, kilobase-length DNA subject to sub-picoNewton mechanical forces. Decondensation by high force proceeds through a series of discrete increases in extension, indicating that PARP1 stabilizes loops of DNA. This model is supported by DNA braiding experiments which show that PARP1 can bind at the intersection of two separate DNA molecules. PARP inhibitors do not affect the level of condensation of undamaged DNA, but act to block condensation reversal for damaged DNA in the presence of NAD^+^. Our findings establish a mechanism for PARP1 in the organization of chromatin structure.

## INTRODUCTION

The poly(ADP-ribose) polymerase (PARP) family of enzymes catalyse the transfer of ADP-ribose units to target proteins. The human genome encodes 17 PARP family genes which play a wide variety of roles in cellular processes^1,2^ with PARP1 accounting for 90% of ADP-ribose synthesis^3^. Since the 1980s, it has been known that the catalytic activity of PARP1 is stimulated by binding at single-strand and double-strand breaks in DNA^4^. This activity is a key stage in single-strand DNA break repair, with the formation of poly(ADP-ribose) acting to loosen chromatin^5^ and to recruit additional DNA repair factors^6,7^. Much recent work on analysing PARP1 has been stimulated by the discovery that inhibition of PARP activity results in synthetic lethality towards BRCA mutated cancer cells^8,9^. PARP inhibitors trap the protein at sites of DNA damage by inhibiting enzymatic activity and thereby block poly(ADP-ribose) induced unbinding^10–13^. This leads to increased double-strand breaks via collapse of replication forks^14,15^ and confers sensitivity of BRCA mutated cells to PARP inhibitor drugs.

PARP1 is a multi-domain protein with three zinc fingers at its N-terminus followed by an automodification domain, a DNA binding WGR-motif domain and a catalytic domain at the C-terminus^16^. The first two zinc fingers coordinate binding at a single-strand DNA break by forming contacts between the exposed bases and hydrophobic residues of the protein^17,18^. By binding at a site of DNA damage, PARP1 initiates the assembly of its multiple domains into a compact structure^19^. This results in the unfolding of specific autoinhibitory structures in the catalytic domain, thereby enabling access of NAD^+^ to the active site and subsequent poly(ADP-ribose) synthesis^20,21^. Biochemical studies have shown that PARP1 has a high affinity for DNA lesions, and a moderate affinity for undamaged DNA. For PARP1 binding to double-strand or single-strand breaks at physiological salt concentrations, typical values for KD are in the range of 10-100 nM^21–23^. The affinity of PARP1 for an 18 bp, ligated, symmetric dumbbell, which mimics undamaged DNA, was measured as K_D_ = 1 μM at 100 mM NaCl concentration17. Studies on the dynamics of PARP1 show that it rapidly accumulates at sites of DNA damage in the nucleus^24^ using intersegmental transfer along DNA to increase the speed of damage localization^25^.

Together with its role in DNA repair, PARP1 acts as a chromatin-architectural protein and transcriptional regulator^26–28^. PARP1 binds tightly to nucleosome-decorated DNA and represses transcription^29,30^. There are on the order of a million PARP1 molecules per cell^31^, which is equivalent to approximately one PARP1 molecule per 20 nucleosomes^22^, making it one of the most abundant nuclear proteins, with a nuclear concentration of ~20μM. This statistic underlines the potential of PARP1 to shape chromatin architecture. In addition to its enzymatic activation by DNA damage, PARP1 can also be activated by various environmental cues such as heat-shock, which causes increased gene expression^32^. Histone modifications are likely activators in this context^33–35^. Overall, the multifunctional roles of PARP1 motivate a need to increase our understanding of its DNA binding modes and the effects of therapeutic drugs on PARP1-DNA interactions.

In this paper, we have used a combination of single-molecule approaches to measure PARP1-DNA structural dynamics in real-time. We find that PARP1 causes compaction of both damaged and undamaged DNA at sub-pN tension and that high forces result in stepwise unbinding of PARP1. We also find that PARP1 can bridge two separate DNA molecules that are brought in close proximity by DNA braiding. These observations lead us to propose a loop-stabilization model where PARP1 can bridge the intersection of two DNA double-strands. The presence of DNA damage and NAD^+^ causes rapid destabilization of DNA loops - an effect which is blocked by catalytic site inhibitors.

## RESULTS

### Fluorescence microscopy demonstrates condensation of DNA by PARP1

PARP1 binding to long stretches of undamaged DNA was visualized via total internal reflection fluorescence (TIRF) video microscopy using sytox-orange labeled λ-DNA (48.5 kbp long). The DNA was bound at one end to the microscope coverslip via a biotin-streptavidin linkage; continuous solution flow, applied with a syringe pump (Figure 1A), stretched-out the molecule to approximately 70% of its ~16 μm contour length. Before PARP1 addition, the DNA exhibited length fluctuations due to thermal forces; after addition of 400 nM PARP1 the DNA condensed towards its attachment point on the surface (Figure 1B and Figure S1 for full field of view). Condensation commenced at the free-end of the molecule, where a compact structure initially formed and then gradually increased in intensity as it moved towards the anchor point (Figure 1C). It is important to note that the hydrodynamic flow stretches DNA molecules with a line-tension that decreases from a maximum value at the molecule’s attachment point to zero at the free end^36,37^. Hence, while our TIRF results clearly demonstrate DNA condensation by PARP1 is initiated at the free end, it is not possible to distinguish whether this is caused by the presence of a double-strand break or the lower tension towards the end.

**Figure 1.**
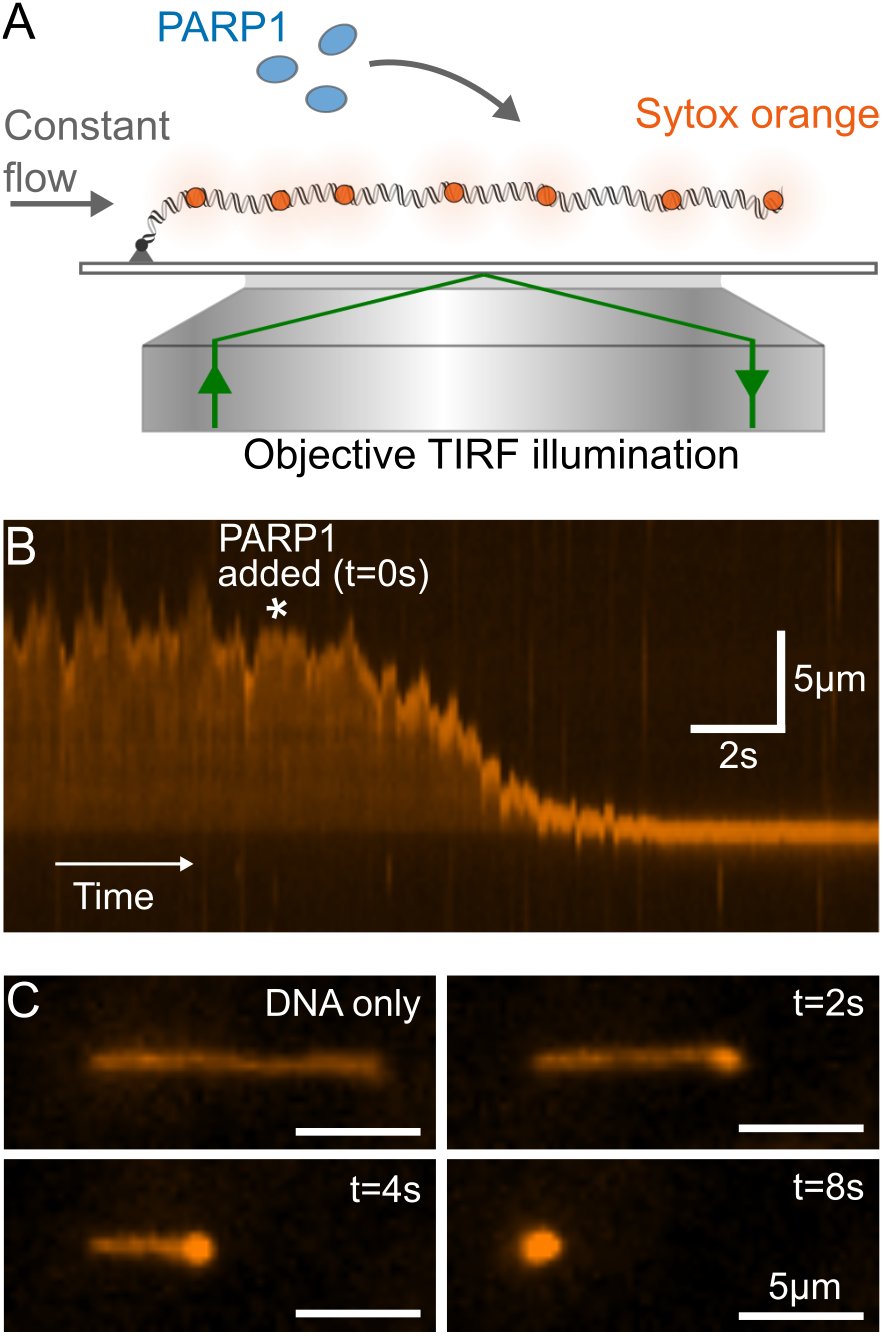
TIRF imaging shows condensation of DNA by PARP1. (A) Schematic of TIRF microscopy imaging of a single λ-DNA molecule stained with Sytox Orange. A constant flow was maintained which stretches out the DNA close to its contour length. (B) Kymograph showing DNA extension over time. 400 nM PARP1 is added at the timepoint indicated by the asterisk. (C) Snapshots showing individual image frames at the indicated timepoints.

### AFM shows decoration of undamaged DNA by single PARP1 molecules

To understand the nature of PARP1 binding to stretches of undamaged DNA, we visualized PARP1-DNA complexes by atomic force microscopy (AFM) in liquid. Firstly, we incubated PARP1 with a 4.4 kbp relaxed, covalently-closed circular plasmid, i.e., an undamaged DNA molecule. The sample was next adsorbed at a mica surface that was functionalized with PLL-PEG, to adhere the DNA while also minimizing non-specific surface adsorption of the PARP1^38^. This resulted in extensive decoration of the plasmids with PARP1 molecules, appearing as bright dots on top of the DNA (Figure 2A), with no apparent preference for any specific location along the DNA. By comparison of multiple images, we found that this decoration did not result in significant compaction of the plasmid probably due to the fact that the extension of DNA in these experiments may largely be determined by its binding to the AFM substrate, trapping it in non-equilibrated, compacted conformations independent of protein binding^39^.

**Figure 2.**
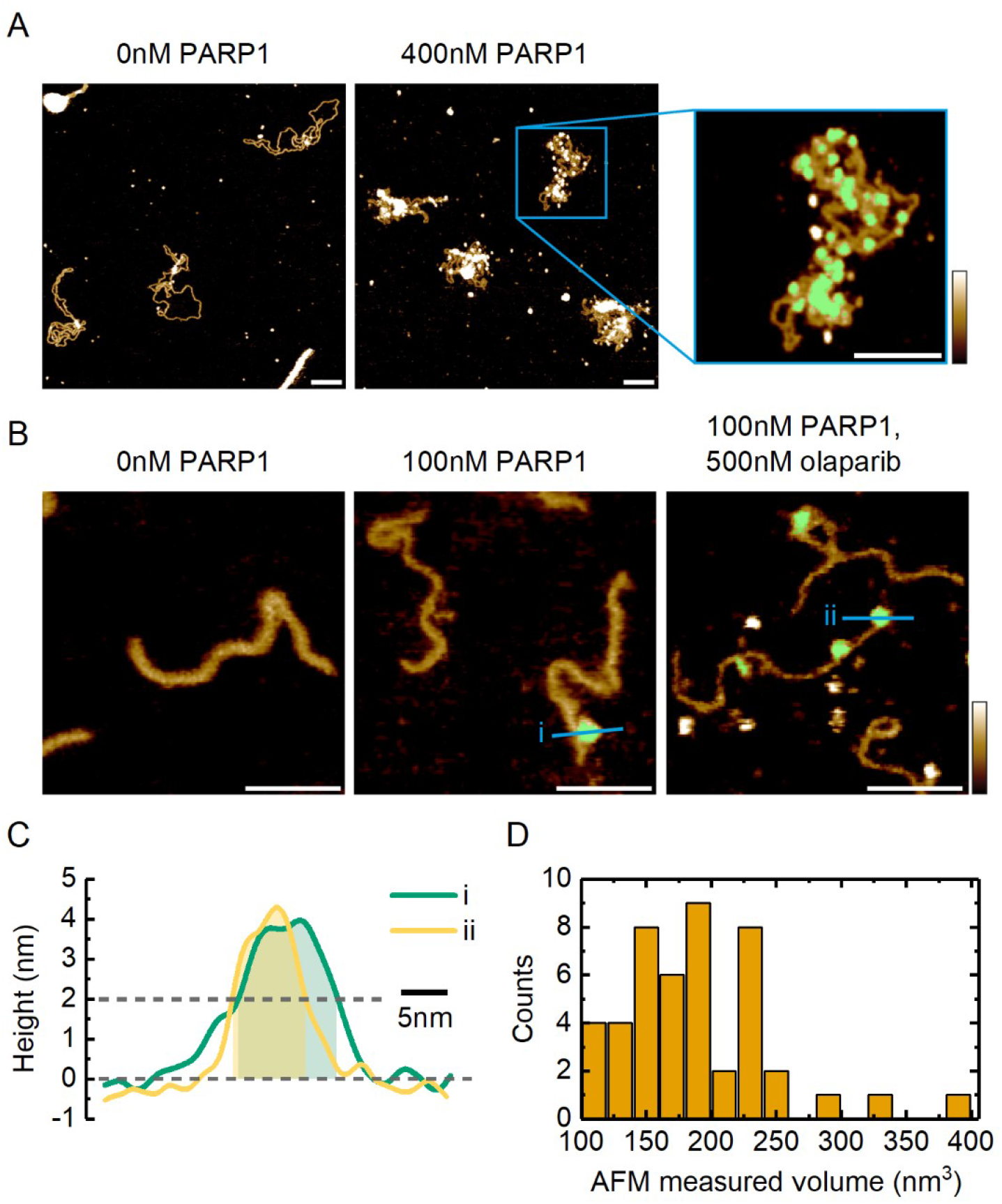
AFM characterization of PARP1-DNA binding. (A) AFM images of PARP1 binding to a 4.4 kbp, covalently closed plasmid. Inset: zoom-in image showing PARP1 bound to plasmid DNA. The presence of bound PARP1 is indicated in green, representing the pixels where the local height exceeded a threshold of ~2 nm above the background. Scale bar = 100 nm. Colour scale = 2.5 nm. (B) Images showing PARP1 binding to a 496 bp PCR fragment with a single-strand break approximately one third of the way along the contour length. The green indicates bound PARP1 that was selected for volume estimation. Scale bar = 50 nm. Colour scale = 2.5 nm. (C) Two selected height profiles with corresponding proteins shown in (B). We observe similar height profiles for PARP1 molecules bound to DNA in the presence and absence of 500nM olaparib. The transparent areas indicate the masked portion of the profiles that contributed to the volume estimate. (D) Histogram of observed PARP1 volumes when bound to the 496 bp PCR fragment.

For comparison, we also imaged PARP1 binding to a short, 496 bp PCR fragment that contained a single-strand break site one third of the way along its contour length, in the absence and presence of a PARP1 inhibitor (olaparib). As shown in Figure 2B, we observed bright dots – attributed to PARP1 – at locations that were consistent with its binding to the single-strand break, but also to undamaged sections of the DNA, in agreement with previous AFM results on dried samples^40,41^. By considering the local height of these dots above the substrate (Figure 2C), we calculated the observed volume distribution (Figure 2D), which had a median of 182 nm^3^. From the crystal structure of PARP1 we estimated an equivalent spherical volume of 150 nm^3^ (Figure S2). Therefore the images of isolated spots are consistent with a monomer of PARP1. This agrees with previous AFM measurements of PARP1^41^, also when taking into account possible effects of the AFM tip on the appearance of the molecule (Figure S3).

This volume measurement also facilitated the quantification of the number of bound PARP1 molecules on the plasmid DNA: To this end, we created a mask that defined areas of PARP1 binding (Figure 2A: magnified section), and then calculated the total volume under the mask before dividing by the volume of a single PARP1 molecule. Analysis of multiple images yielded an estimate of the number of bound PARP1 molecules to be 16 ± 3 (mean ± s.e., N = 18) at 200 nM PARP1 and 59 ± 15 (mean ± s.e., N = 13) at 400 nM PARP1 (Figure S4). Finally, to verify the robustness of our observations against variations in sample preparation, we also analysed DNA that was exposed to PARP1 after DNA adsorption at the AFM substrate. While this resulted in lower amounts of bound PARP1, it yielded qualitatively similar results (Figures S5-S6).

### Real-time measurement of DNA condensation by magnetic tweezers

Noting the PARP1-induced condensation of DNA as observed by TIRF microscopy and the overall decoration of undamaged DNA by bound PARP1 molecules, we next sought to characterize the PARP1-induced condensation with precise control of tension and DNA damage, using magnetic tweezers. We tethered individual DNA molecules (7.9 kbp in length) to a magnetic bead and applied controlled forces and rotations via a pair of permanent magnets (Figures 3A, B). The DNA extension was measured by real-time video tracking of the bead position, while the force was controlled by varying the separation between the bead and the magnets. Importantly, the use of magnetic tweezers enabled us to verify that there was a single, undamaged DNA molecule tethered between the surface and bead, based on the characteristic change in DNA extension upon bead rotation at low force^42^ (Figure 3C). In addition, as the applied tension (force) was reduced from 5.8 to 0.07 pN (Figure 3D), DNA extension in the absence of PARP1 decreased in a manner consistent with the worm-like-chain force-extension property, typical for single DNA molecules (see Figure 4a)^43^.

**Figure 3.**
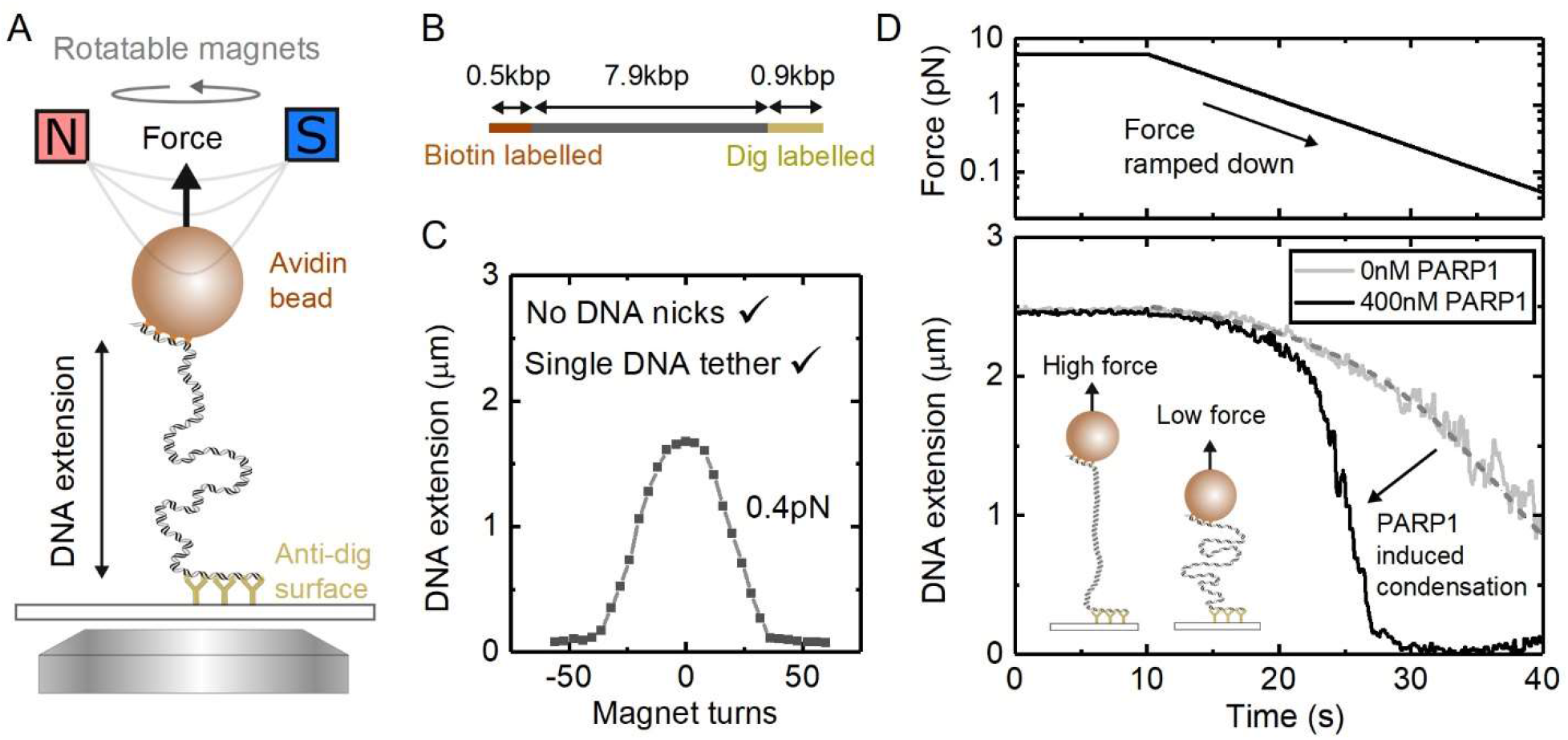
Magnetic tweezers showing PARP1 induced condensation on undamaged DNA. (A) Schematic of magnetic tweezers showing DNA stretched between the bead and coverslip. The height of the magnets controls the force acting on the bead and the bead can be rotated by rotating the magnets. (B) DNA construct used for magnetic tweezers with two labelled handles acting as attachment points. (C) Single DNA tethers without nicks are selected by examining the characteristic response of DNA extension to magnet turns. (D) Effect of force ramp on DNA extension in the presence of 400 nM PARP1 and 0 nM PARP1. At low forces (<1 pN), PARP1 induces significant compaction of the DNA. The dashed grey line shows a fit of the worm-like chain model to the DNA extension for 0 nM PARP1 with a fixed persistence length of 50 nm. The DNA contour length was a free parameter for the fit and yielded 2.7 μm in good agreement with the expected length of 2.6 μm for the 7.9 kbp DNA.

**Figure 4.**
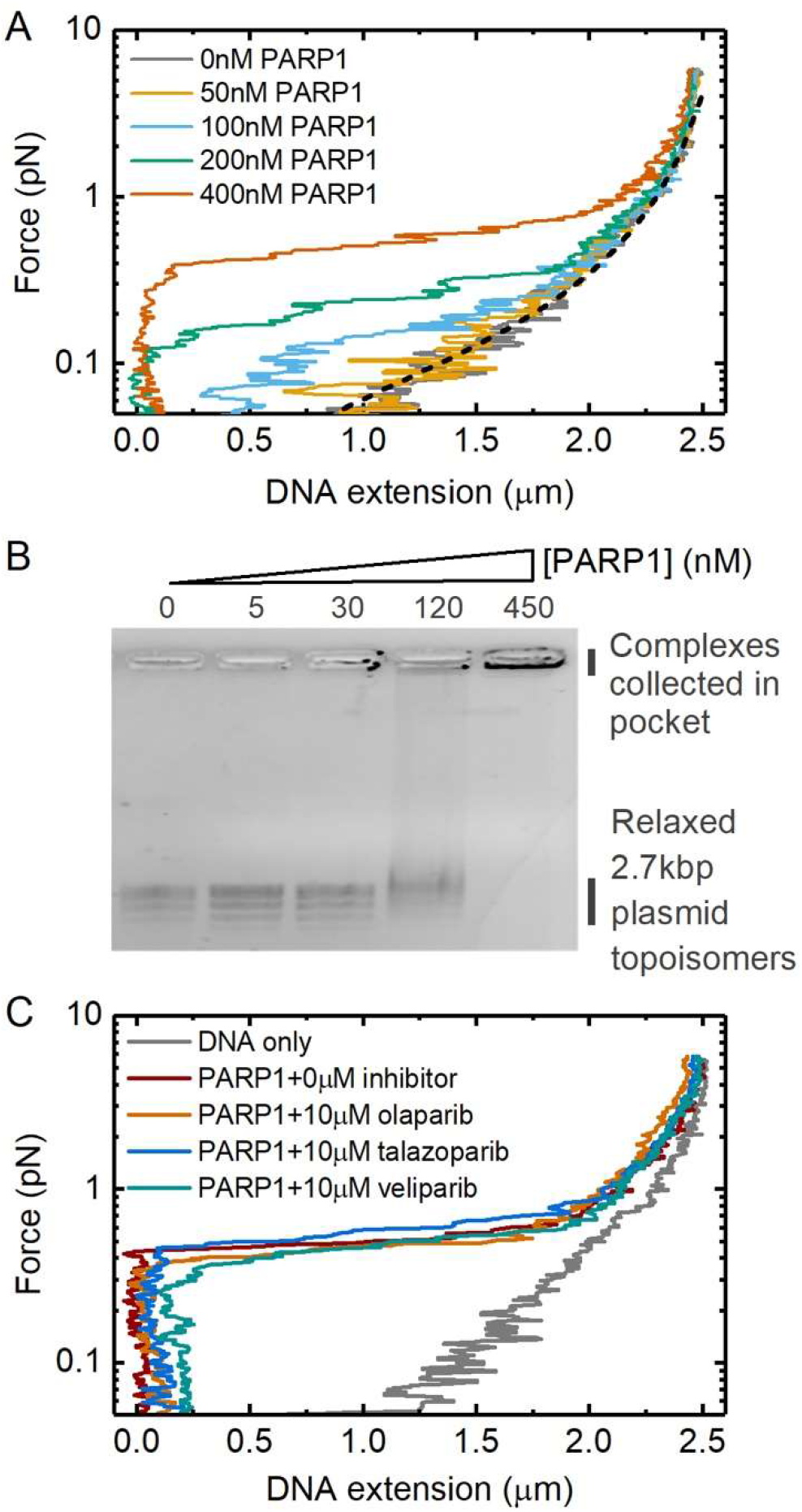
Concentration dependence of DNA condensation by PARP1. (A) Magnetic tweezers data showing the change in DNA extension at different applied forces. The dashed black line indicates the worm-like chain model prediction for naked DNA. (B) Agarose gel electrophoresis showing titration of PARP1 with 3 nM of relaxed, covalently closed 2.7 kbp DNA plasmid. Several bands are visible for the plasmid sample, corresponding to different topoisomers. (C) Force-extension curves measured in the presence and absence of three PARP inhibitors. 400 nM PARP1 concentration was used in each case.

While the addition of 400 nM PARP1 made little difference to DNA length measured at higher forces (Figure 3D), it caused a dramatic shortening of the DNA as force was reduced below 1 pN, consistent with our observation by TIRF microscopy. To confirm that the observed compaction was specific to PARP1 and not due to any co-purifying contaminants, we performed a control experiment that showed no DNA compaction when PARP1 was specifically depleted by immunoprecipitation (Figure S7). We also repeated the experiment using the same (48.5 kbp) λ-DNA template, as used in our previous TIRF experiments, and confirmed the same force-extension behavior, with DNA compaction occurring at forces below 1 pN in the presence of 400nM PARP1 (Figure S8). Taking these results together, we conclude that PARP1-induced condensation does not depend on the presence of a free end or double-strand break (as in the TIRF experiments in Figure 1), but does strongly depend on the DNA tension.

The condensation of undamaged DNA also showed a strong dependence on PARP1 concentration, as measured via force-extension curves at different PARP1 concentrations (Figure 4A). The force was ramped using the same profile as shown in Figure 2D. DNA condensation was observed at concentrations of 100 nM PARP1 and above, and for increasing PARP1 concentration, higher forces were required to extend the DNA to lengths observed in the absence of PARP1. To characterise the condensation at effectively zero force, we analysed binding to relaxed, covalently-closed plasmid DNA by agarose gel electrophoresis. This showed the formation of PARP1-DNA complexes at 120 nM PARP1 and above, as indicated by a smearing of the DNA plasmid bands (Figure 4B). At 450 nM PARP1 concentration, the complexes did not run through the gel and collected in the gel pocket. We also measured the effect of several PARP inhibitors on the condensation of undamaged DNA (Figure 4C). At 10 μM concentration of each of olaparib, talazoparib and veliparib, we observed no effect on PARP1-induced DNA condensation. For comparison, experiments with PARP2 showed no observable condensation at concentrations up to 1 μM (Figure S9).

### Magnetic tweezers reveal stepwise unbinding and DNA bridging by PARP1

To gain further insight into the mechanism of PARP-induced DNA condensation, we examined the reversal of condensation by monitoring the time-course of DNA extension changes following application of a rapid increase in force applied to the magnetic bead. Figure 5A shows an example trace where we applied a step-change in force from 0.6 pN to 2.2 pN and measured the resulting response in DNA extension over time. In the absence of PARP1, a step-change in force produces a rapid change in DNA extension. In the presence of PARP1, however, we observed a different behaviour. Specifically, DNA extension showed initially a rapid change in length followed by a series of discrete steps to reach its final extension (marked by arrows in the lower panel of Figure 5A). We collected 44 traces and used a chi-squared minimization method to detect the time and amplitude of the stepwise length changes following the force-step^44,45^. Figure 5B shows a histogram of the measured step amplitudes, with an average step size of 158 ± 10 nm (mean ± s.e.).

**Figure 5.**
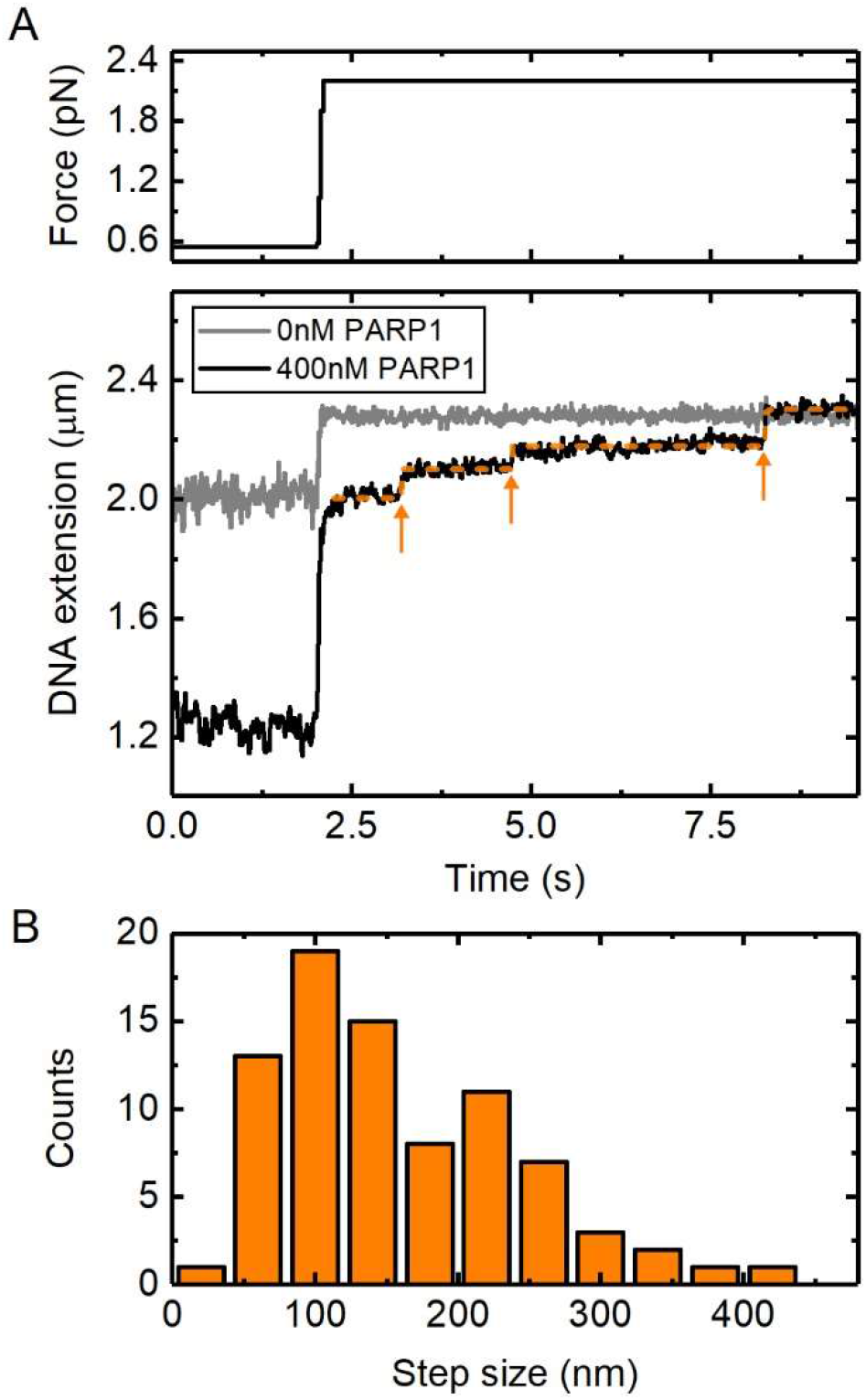
Stepwise reversal of PARP1-induced condensation, as observed upon application of high forces. (A) Traces for 0 nM PARP1 and 400 nM PARP1, showing the changes in DNA extension after a step change in force from 0.6 pN to 2.2 pN. The orange line indicates the result of the step-fitting algorithm used with the arrows showing the positions of individual detected steps. A delay of 0.2 s between the force change and the beginning of step detection was imposed. (B) Histogram of the measured step sizes. 81 steps were detected from a total of 44 traces where the force was stepped from 0.6 pN to 2.2 pN.

The observed step sizes are 2-5 times larger than the approximate circumference of PARP1 (see Figure S2), arguing against a model where DNA is tightly wrapped around PARP1, such as in a nucleosome. Instead, the large step amplitude, together with the sub-1 pN threshold for condensation (Figure 4A), suggests that PARP1 stabilizes loops of DNA that are formed under thermal fluctuations. The statistical mechanics of DNA under tension shows that DNA loops, formed by thermal fluctuations, are strongly suppressed when the force applied to DNA is greater than 1 pN^46,47^. This sub-pN force threshold is in agreement with other proteins which are known to stabilize DNA loops^48,49^. Worm-like chain models of DNA force-extension behaviour indicate the expected distribution of loop sizes will be centred at several times the 50 nm persistence length of DNA with a range of several hundred nanometres^47^. The loop-size distribution is due to a competition between the enthalpic cost of forming tight DNA bends, disfavouring short loops and the entropic cost of bringing two distant regions of DNA close enough to intersect which therefore disfavours long loops.

A DNA looping mechanism requires that two sections of DNA are bridged where they intersect. To test whether such a mechanism might apply to PARP1-induced DNA condensation, we investigated the ability of PARP1 to bind at the intersection of two dsDNA strands. In these experiments, the surface density of DNA on the magnetic beads was increased by incubation with a higher concentration of DNA. This resulted in a significant number of beads that were connected to the microscope coverslip by two DNA molecules. A single rotation of such, doubly-tethered, beads causes a braid to form between the two DNA molecules, which leads to a dramatic decrease in DNA extension, as observed via the decrease in bead height (Figure 6A)^50,51^. In a typical field of view, the majority of beads were still bound via a single DNA tether giving confidence that attachment via three or more tethers was rare. The size of the height decrease resulting from a single bead rotation is dependent on the exact position of the anchoring points of the two tethers on the bead and surface. In the absence of protein, braid formation is reversible and can be unwound simply by rotating the bead back to its starting position. Figure 6B presents data traces acquired using three independent beads imaged in the same experiment. Each bead showed the characteristic change in extension upon rotation in the absence of PARP1. In the presence of 200 nM PARP1, we observed that the extension was stabilised at the braided level when the magnets are first rotated by one turn to form the braid and then rotated back to the start position (Figure 6C). This shows that PARP1 bridges the two DNA molecules at their crossover point and prevents them from coming apart when the two strands are unwound. After flowing through buffer to remove free PARP1 from the flow-cell (Figure 6D), we observe rapid unbinding of the PARP1 as shown by the unwinding of the DNA braids in less than 30s.

**Figure 6.**
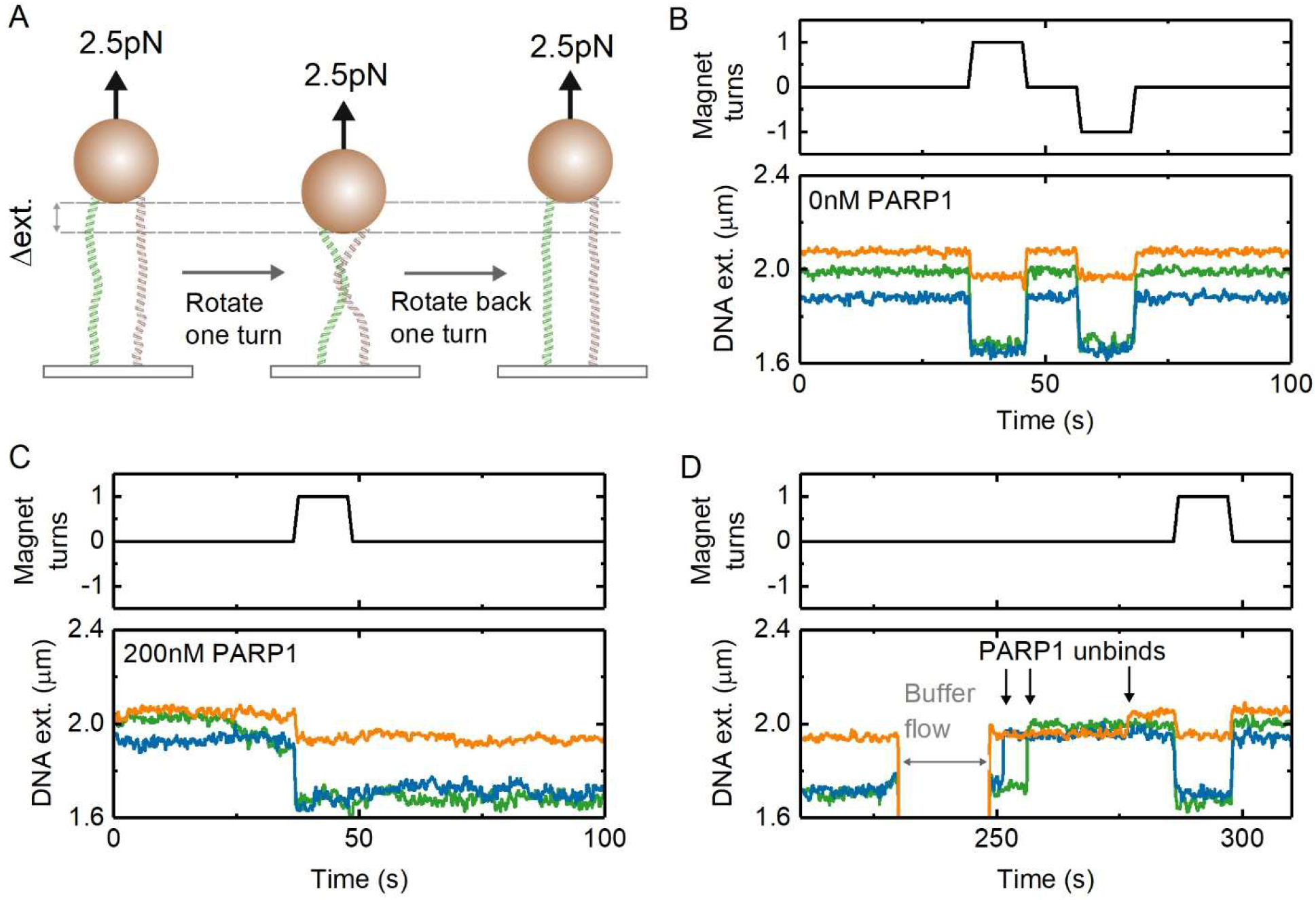
Bridging of two DNA molecules by PARP1. (A) Schematic of experiment – rotating a dual tethered magnetic bead results in the formation of a DNA braid and a change in measured extension. (B) In the absence of PARP1, rotation of the magnetic bead clockwise (+1 turns) or counter-clockwise (−1 turns) results in a reversible change in DNA extension. The three colours represent data recorded simultaneously from three beads in the field of view. (C) In the presence of 200 nM PARP1, the effect of a one-turn rotation is found to be irreversible, until (D) the flow-through of 0 nM PARP1 buffer (at t = 230 s) results in a rapid reversal to original DNA extension. Arrows indicate steps where DNA extension returns to original value.

### Kinetics of condensation loss in the presence of DNA damage and NAD^+^

So far, we have focussed on the effect of PARP1 binding to undamaged DNA. However, as noted in the introduction, PARP1 is mostly known for its role in DNA damage repair, a process initiated by the synthesis of strongly negatively charged poly(ADP-ribose) chains (PAR) at sites of DNA damage. The PAR chains are covalently coupled both to PARP1 itself (auto-modification) and to other nearby proteins. As the auto-modified PAR chains grow in size, they are thought to destabilise PARP1-DNA interaction and cause PARP1 to release from DNA^12^. To investigate this process and its effects on DNA condensation, we extended our investigations to DNA that presented single-strand breaks (SSBs), introduced by site-specific nicking endonucleases: Nt.BsmAI and Nb.BsmI. Single DNA tethers were selected and 14 SSBs were made in the 7.9 kbp DNA construct (Figure 7A). The formation of SSBs was monitored *in situ* by first checking the topological continuity of the dsDNA tethers by measuring their characteristic shortening in response to supercoiling and plectoneme formation. Then, following endonuclease addition, formation of single-strand nicks (SSBs) allowed the DNA to rapidly relax back to its length at zero turns since the plectonemes unwound via free rotation at the nick sites (Figure 7B).

**Figure 7.**
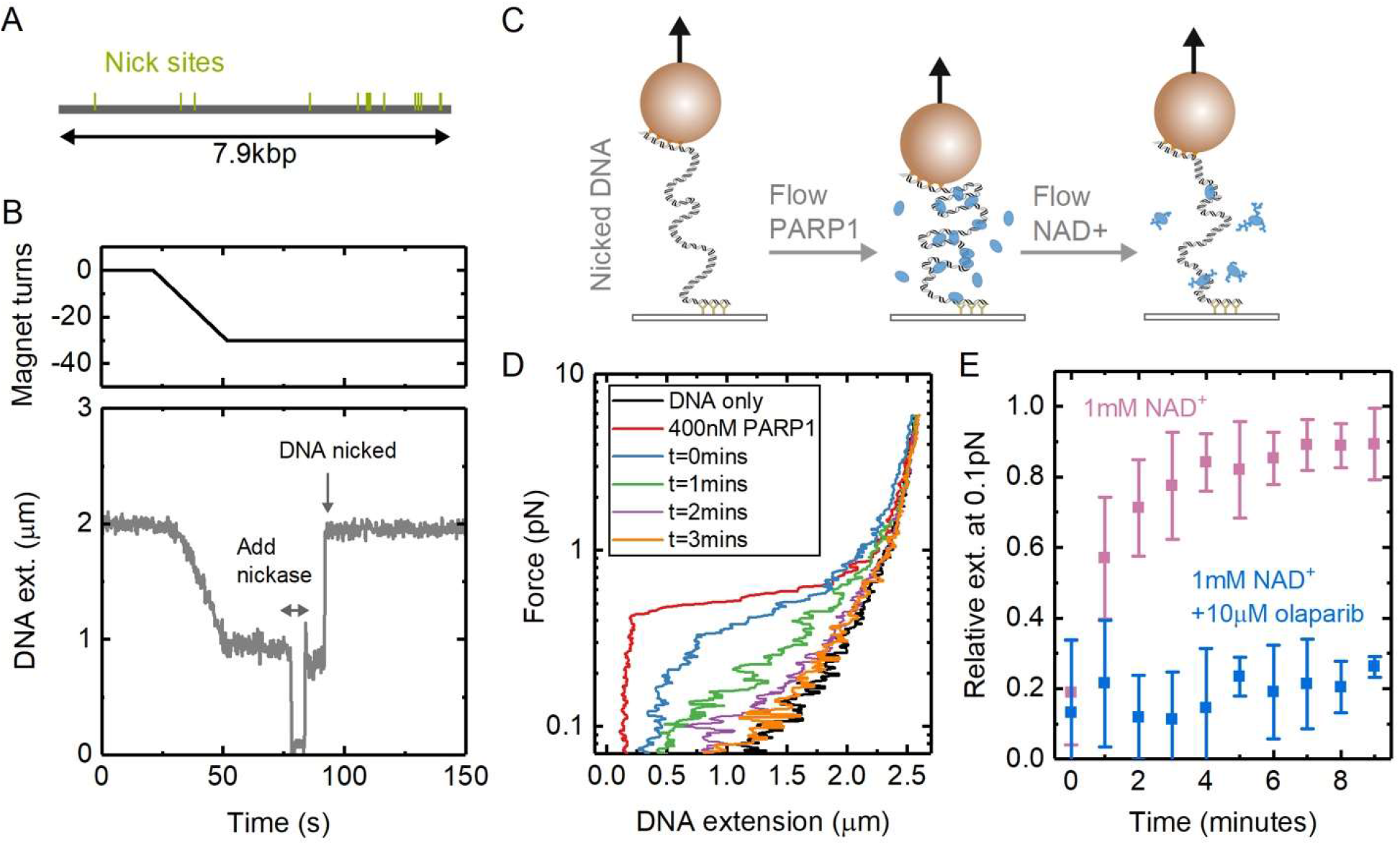
Reversal of PARP1 induced DNA condensation in the presence of NAD^+^ and DNA damage. (A) Positions of 14 nick sites on 7.9 kbp DNA section of magnetic tweezers for the nicking endonucleases Nt.BsmAI and Nb.BsmI. (B) Example trace showing the formation of nicks visualised in real-time with magnetic tweezers. The DNA is coiled by magnet rotation before addition of nicking endonuclease which results in rapid uncoiling. (C) Schematic of experiments for measuring the time dependence of condensation. PARP1 is added to damaged DNA before flowing through solution containing NAD^+^. (D) Force-extension curves showing the change in condensation after adding 400nM PARP1 followed by adding 1 mM NAD^+^. (E) Extension, relative to naked DNA, measured at 0.1 pN force as a function of time after flowing through NAD^+^ containing solution. The graph shows results from two experiments 1) addition of 1mM NAD^+^ (N = 9 magnetic beads) and 2) addition of 1mM NAD^+^ + 10μM olaparib (N = 4). The error bars show standard deviation.

After creation of the single-stranded breaks, we added 400 nM PARP1 to the solution and measured DNA condensation from the force-extension relationship (Figure 7C,D). This resulted in similar behaviour as observed for the undamaged DNA (cf. Figure 4A). We next flushed the flow cell with a solution containing NAD^+^ and measured how condensation varied over time by repeating the force-extension curve every minute. Figure 7D shows this change for a buffer solution without PARP1 but containing 1 mM NAD^+^. In Figure 7E, we quantify the change in condensation by plotting the extension relative to naked DNA at 0.1 pN as a function of time after adding the NAD^+^ containing solution. We found >70% recovery (i.e. loss of condensation) within 3 minutes of adding NAD^+^. Release of PARP1 from the DNA and reduction in the condensation effect can be explained by formation of poly(ADP-ribose) chains which are known to reduce PARP1-DNA affinity. We then tested the effect of adding 1 mM NAD^+^ in the presence of 10 μM olaparib and found that DNA remained in its condensed state for the duration of the ten-minute experiment and no longer returned to the force-extension behaviour characteristic of naked DNA. This behaviour is consistent with olaparib competing with NAD^+^, blocking poly(ADP-ribose) formation and preventing PARP1 from unbinding the DNA. For undamaged DNA we found a recovery of >70% extension over eight minutes when buffer not containing NAD^+^ was flowed through (Figure S10).

## DISCUSSION

We have used a combination of single-molecule techniques to show that PARP1 condenses both damaged and undamaged DNA. The condensation is reversible by applying a high force (>1 pN). The loss of condensation from application of a step change to high force was observed to occur in a series of discrete changes in extension. The relatively large amplitude of the steps (>100 nm) is not consistent with DNA wrapping around PARP1 but is more easily explained by formation of DNA loops. The wide distribution of step-size amplitudes is also consistent with initial formation by thermal fluctuations and capture by PARP1 binding^47^. The ~1 pN threshold of condensation is also typical of DNA looping seen in previous studies^46^. To test the idea that PARP1 can stabilise DNA-cross-over points we also performed DNA braiding experiments. These experiments clearly showed that PARP1 forms a bridge between DNA molecules, stabilizing the DNA-DNA intersection as required for a DNA looping activity. Our AFM investigation confirmed that PARP1 binds to undamaged plasmid DNA forming a disordered structure although individual looping events were not resolved, possibly because of destabilisation due to the binding to the AFM substrate. Figure 8 depicts our model for the DNA condensation activity of PARP1 on a single molecule of double-stranded DNA. Thermal fluctuations create intersections of DNA when the tension in the DNA is sufficiently low (below ~1 pN). PARP1 bridges and stabilises these thermally induced loops thereby causing a reduction in DNA extension. For single DNA molecules, DNA loop-stabilisation shows a strong dependence on DNA tension since the probability of loop formation is highly force dependent^52,53^.

**Figure 8.**
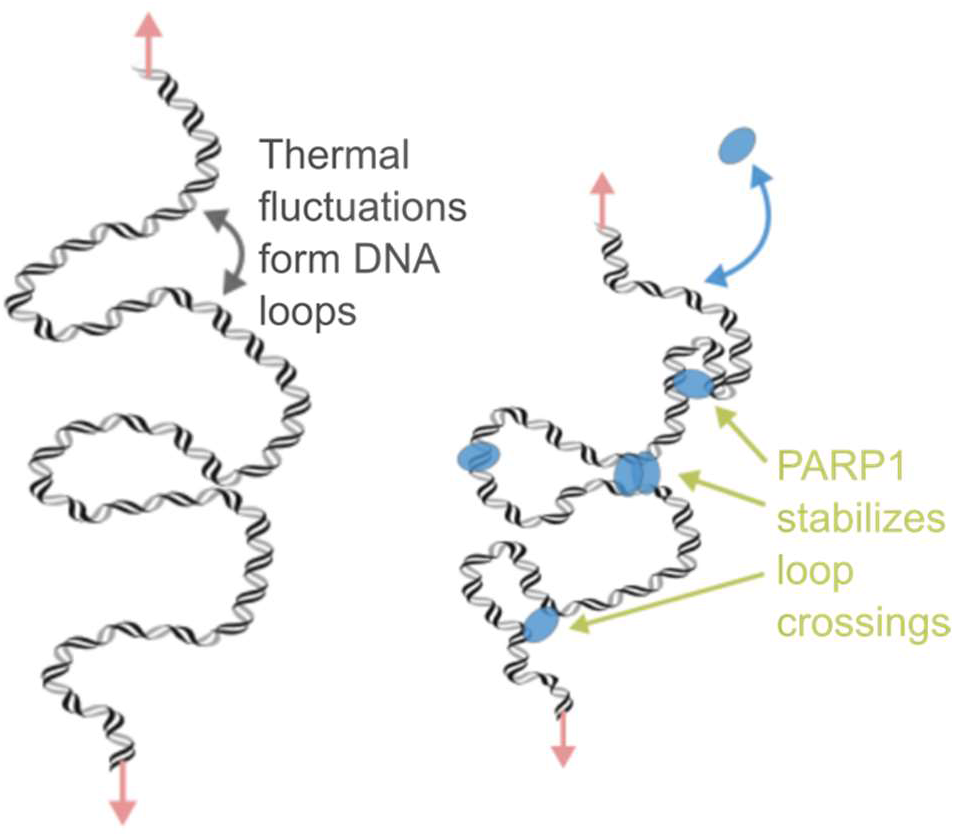
Loop stabilization model for PARP1 condensation of DNA. Left: In the absence of protein and at low forces, thermal fluctuations induce the formation of DNA loops. Right: The multiple DNA binding sites of PARP1 enable bridging to stabilize the loop and thereby reduce the DNA extension.

The loop-stabilisation activity of PARP1 is consistent with recent results that show that PARP1 can move through the genome using intersegmental transfer^25^, since both observations require the use of multiple DNA binding sites. Moreover, our measurement of fast unbinding from DNA braids (Figure 6D) supports the idea of rapid exchange of PARP1 at intersections between multiple DNA strands. Our experiments, however, do not directly determine the number of PARP1 molecules needed to bridge a single DNA loop. While the crystal structure of PARP1 shows that there are multiple DNA contacts formed through the zinc fingers and WGR domain^19^, dimers or oligomers of PARP1 could also be formed at the points of loop-stabilisation. Further research using PARP variants and mutants will be of interest to pin down the necessary domains for DNA condensation and therefore the likely number of molecules forming DNA bridges. Towards this goal, we measured force-extension curves in the presence of 1 μM PARP2 which showed no effect on DNA extension. PARP2 lacks the Zn finger domains of PARP1 thereby suggesting that these domains are important in mediating loop-stabilization.

PARP inhibitors are known to trap PARP1 in an insoluble fraction of chromatin in cellular assays^10^. The mechanism behind the level of trapping for different inhibitors has been investigated with various in vitro biochemical methods using damaged DNA^11,13^. Since short DNA molecules are used in most binding studies such as fluorescence polarization or surface plasmon resonance, these unavoidably include DNA hairpins or ends preventing the unambiguous measurement of PARP1 binding to undamaged DNA. Our magnetic tweezers assay allows us to measure real-time PARP1 binding to undamaged DNA since the free ends are outside the measurement region. The results did not show any effect of high concentration of inhibitors on PARP1 condensation of undamaged DNA. When we introduced damage to the DNA through single-stranded breaks, we found that the inhibitor olaparib slowed the rate of NAD^+^ induced decondensation. It is known that formation of PAR chains reduces the affinity of PARP1 for DNA breaks by inducing strong electrostatic repulsion^12^. Our results are consistent with this effect and show that PARylation also affects the ability of PARP1 to condense DNA.

In conclusion, using single molecule techniques, we have shown that PARP1 condenses DNA by a loop-stabilisation mechanism. PARP1 is known to play an important role as a chromatin architectural protein together with its role in DNA repair^54^. Independent of its PARylation activity, it has been observed that PARP1 condenses nucleosome-bound DNA^55^ and is enriched at regions of heterochromatin^56,57^. In *Drosophila*, PARP1 is also known to inactivate certain transcriptional domains and repress retrotransposable elements^58,59^. Loop-stabilisation is a novel mechanism which can explain the fundamental physical interaction between DNA and PARP1 that underlies these condensed areas of chromatin. Methods to probe cellular chromatin architecture and protein induced looping^60,61^ should provide further insight into the importance of DNA looping by PARP1 in the context of the nucleus.

## METHODS

All reagents were from Sigma-Aldrich UK unless otherwise stated.

### Total Internal Reflection Fluorescence (TIRF) microscopy

λ-DNA (NEB) was functionalized at one end by annealing with a biotinylated oligonucleotide/5Phos/AGGTCGCCGCCCTTTTT/Bio (IDT) followed by ligation with T4 DNA ligase (NEB). The DNA was purified using a bead purification kit (Qiaex II, Qiagen). A flow-cell was constructed using polydimethylsiloxane (Sylgard 184, Dow), patterned double-sided tape (AR90880, Adhesive Research) and a glass coverslip (Menzel) coated with 2% nitrocellulose in amyl acetate. The flow cell was incubated for 10 minutes with 1 mg/ml streptavidin (NEB) and then 10 minutes with Blockaid (ThermoFisher). Biotinylated λ-DNA was then injected and allowed to bind to the streptavidin coated surface. Imaging was performed using a buffer of 20 mM HEPES pH 7.8, 150 mM NaCl, 2 mM MgCl_2_, 0.5 mM TCEP, 1 mg/ml BSA, 1 mg/ml β-casein, 500 nM Sytox Orange (ThermoFisher). A constant flow-rate of 100 nl/s was applied using a syringe pump (Harvard Instruments PhD 2000). The sample was illuminated by a 532 nm laser in objective TIRF mode with a 60x Nikon objective and images acquired at 10 fps with an EMCCD camera (Andor iXon).

### Atomic force microscopy (AFM)

A 496bp section of λ-DNA was amplified by PCR using Taq polymerase, forward primer 5’- TGAAATTGCCGCGTATTACGC-3’ and reverse primer 5’-TTTCTCGTAGGTACTCAGTCCG-3’. The PCR product was purified by a kit (QIAquick, Qiagen). The sequence of the 496 bp product contains a site for Nt.BsmAI between bases 172 and 173, i.e., approximately one third of the total length. Nicking was performed by incubating the DNA with Nt.BsmAI and subsequently the DNA was further purified (QIAquick, Qiagen). Relaxed pBR322 plasmid DNA was purchased from Inspiralis Ltd. The block copolymer mPEG_5k_-b-PLKC_10_, methoxy-poly(ethylene glycol)-block-poly(L-lysine hydrochloride), was purchased as a lyophilized powder from Alamanda polymers. PLL_150-300k_ (poly-L-lysine, 0.1% w/v, 150,000-300,000 MW) was purchased from Sigma-Aldrich.

For AFM imaging, a freshly cleaved mica specimen disk (diameter 6 mm, Agar Scientific, UK) was functionalized with PLL-PEG, consisting of a mixture of mPEG_5k_-b-PLKC_10_ (1 mg/ml in Milli-Q water) and PLL_150-300k_, prepared as previously described^38^. The washing and imaging buffer used throughout was 12.5 mM NaCl, 12.5 mM HEPES, 0.5 mM TCEP, pH 7.8, filtered by passage through 0.2 μm syringe followed by a 10 kDa cutoff centrifugal filter (Amicon Ultra, Millipore). Once the modified mica was prepared, 20 μl of linear 496 bp DNA (1.5 ng/μl) or relaxed pBR plasmid DNA (2.5 ng/μl) was then added to the disk and gently mixed. After a 30 minute adsorption, the sample was then washed 5 times and topped up to 30 μl with imaging buffer. For the pre-incubation assays, the DNA was incubated with PARP1 (and 500 nM olaparib if stated) for 30 minutes at room temperature prior to deposition onto the modified mica surface. Images of PARP1 pre-incubated with DNA prior to deposition are shown in Figures 2A-B, S4C and S5C. For the post-incubation assays, after DNA was immobilized and imaged on the modified mica surface, the sample was exposed to the imaging buffer with PARP1 (and 500 nM olaparib if stated). Imaging was resumed after 5 minutes incubation without washing. See Figures S4A and S5A for images. Despite the differences in the quantity of PARP1 bound to the DNA across these two sample preparation methods, a similar qualitative behaviour of PARP1 binding is observed.

All AFM measurements were carried out in buffer and at room temperature. Data were recorded using a Dimension FastScan Bio AFM (Bruker, Santa Barbara, USA), using force-distance-curve based imaging (Bruker’s PeakForce Tapping mode). Force-distance curves were recorded over 10-40 nm (PeakForce Tapping amplitude of 5-20 nm) at a frequency of 8 kHz. FastScan D (Bruker) cantilevers were used for all imaging (nominal spring constant ~0.25 Nm^−1^). Imaging was carried out using PeakForce setpoints in the range of 15-30 mV, with a deflection sensitivity of approximately 18 nm/V. Images were recorded at a scan size of 1-2 μm, with 1024 pixels per line and at a line rate of 4 Hz. Images were processed using either Gwyddion^62^ or a Python that employs the Gwyddion ‘pygwy’ module, as described in greater detail elsewhere^63^. All images were initially subjected to correction by first-order line-by-line flattening, median line-by-line flattening and zeroth-order plane fitting to remove background offset and tilt. The background was also zeroed by setting the mean value to zero. A 1-2 pixel (~1-2 nm) gaussian filter was then applied to remove high frequency noise.

To identify DNA molecules, the Python script was employed to find grains that have heights within 1σ of the mean of the height distribution of the image, and have a pixel area within 50-150% of the median grain area, with the DNA identification verified by direct inspection of the images. The thus identified “grains” were interrogated further to measure the size of PARP molecule(s) bound to linear DNA and the total PARP volume bound per plasmid DNA molecule. To isolate protein molecules bound to DNA, these grains were manually thresholded with a height of of ~1.5x the height of a DNA molecule, equivalent to around 2 nm above the background. Volumes were evaluated as the zero-basis volume in Gwyddion, i.e., the volume from zero height covered by the mask. The images used to measure the size of PARP molecule(s) bound to linear DNA were taken using both sample preparation methods described above, where PARP1 is either pre- or post- incubated with DNA, in the presence or absence of olaparib. The distribution of measured PARP1 volumes bound to linear DNA are visualized as a histogram (Figure 2D).

### Magnetic tweezers DNA construct

The central part of the DNA construct used for magnetic tweezers was a 7.9 kbp fragment produced by restriction digestion of λ-DNA using SapI and BsaI (NEB) and subsequent agarose gel purification. A 478 bp fragment of biotin labelled DNA was produced by PCR with Taq Polymerase (NEB) of λ-DNA using the following primers: 5’-CGAACTCTTCAAATTCTTCTTCCA-3’ and 5’-GATTGCTCTTCTGTAAGGTTTTG-3’ with a 5:1 ratio of dTTP:biotin-11-dUTP (Jena Bioscience). Similarly, an 878 bp fragment of digoxigenin labelled DNA was produced by PCR of λ-DNA with primers: 5’-<u>TAGTCCAGAACGAGACC</u>GCAACAGCACAACCCAAACTG-3’ and 5’-AATCTGCTGCAATGCCACAG-3’ (underline section indicates overhang) with a 10:1 ratio of dTTP:digoxigenin-11-dUTP (Jena Bioscience). The labelled fragments were purified using a PCR purification kit (QIAquick, Qiagen). The 478 bp fragment was digested with SapI and the 878 bp fragment was digested with BsaI before each fragment was again purified. The central fragment and two labelled fragments were then ligated by incubation with T4 DNA ligase (NEB) overnight. The construct was finally purified by agarose gel electrophoresis. All DNA gel extractions were performed without exposure to UV or intercalating dyes by staining the outer two lanes only to calculate the band position and using these as alignments for gel excision. The DNA construct length was confirmed by gel electrophoresis (Figure S11). The DNA was conjugated to streptavidin-coated magnetic beads (MyOne T1 Dynabeads, Thermo Fisher) by incubating for 15 minutes at room temperature. The concentration of DNA required was empirically determined in order to maximize the number of single or double tethers.

### Magnetic tweezers measurements

1.5 μm diameter silica microspheres (Bangs Labs) were suspended in 2% collodion at approximately 0.05 mg/ml. The solution was then spin coated onto a glass coverslip to form a thin layer of nitrocellulose with the embedded silica microspheres acting as fiducial markers for drift tracking. A flow-cell was formed by cutting channels into double-sided tape (AR90445, Adhesive Research) and sandwiching this between the nitrocellulose coated coverslip and a second coverslip. A 0.1 mg/ml solution of anti-digoxigenin antibodies (Sigma-Aldrich) was incubated in the flow cell for 30 minutes followed by surface passivation for 30 minutes with Blockaid solution (Thermofisher). After blocking, DNA labelled magnetic beads were flowed through the chamber and incubated in the flow cell for 10 minutes. Finally, measurement buffer was perfused through the chamber. All measurements were performed in a buffer of 20 mM HEPES pH 7.8, 150 mM NaCl, 2 mM MgCl_2_, 0.5 mM TCEP, 1 mg/ml BSA, 1 mg/ml β-casein. Solution exchanges were performed with a custom-built gravity perfusion system at a flow rate of 500 nl/s. To add single-stranded breaks, Nt.BsmAI (0.1 U/μl) in 1x Cutsmart buffer (NEB) was added to the flow cell and incubated for 5 minutes followed by Nb.BsmI (0.2 U/μl) in 1x NEB 3.1 buffer (NEB) which was also incubated for 5 minutes. The flow cell was then flushed with 20 mM HEPES pH 7.8, 1 M NaCl, 0.1 mg/ml BSA before adding the measurement buffer. PARP inhibitors were purchased from Cayman Chemical and diluted in DMSO. Inhibitor experiments used a final concentration of 1% DMSO.

A bright-field microscope combining a high-power LED (M660L4-C4, Thor Labs), 60x air objective (Nikon) and camera (DMK 33GX249, Imaging Source) were used to image the magnetic beads. A look-up table for calculating z-position was created by stepping the objective using a piezo actuator (P721 PiFoc, Physik Instrumente). Bead positions were tracked, in real-time, using a previously described algorithm^64^. The force, F, was calibrated according to the formula *F*=*k*_*b*_*Tl*/<*x*^*2*^> where *k*_*b*_ is Boltzmann constant, *T* is absolute temperature, *l* is DNA extension and <*x*^*2*^> is variance in bead x-position (parallel to the magnetic field lines). A single-exponential was used to fit the force versus magnet height relationship. A fast frame exposure time of 0.7 ms was used so that image blurring, due to bead motion, was negligible at the forces used in our experiments^65^. Images were acquired at 20 fps for all experiments except for PARP1 unbinding traces (Figure 5) where a frame rate of 200 fps was used. A rank 2 median filter was used for display. Forces were applied to the magnetic beads using a pair of 5 mm cube neodymium magnets (Supermagnete) mounted on a custom-built z-translation stage. A stepper motor coupled to a drive-belt enabled rotation of the two magnets at a speed of one revolution per second.

### Electromobility gel shift assay

pUC19 plasmid (NEB) was relaxed from its supercoiled form by incubating with Topo I (NEB) and purified (QIAquick, Qiagen). The relaxed plasmid was then incubated with PARP1 for 10 minutes in a buffer containing 20 mM HEPES pH 7.8, 150 mM NaCl, 2 mM MgCl_2_, 0.5 mM TCEP, 0.05 mg/ml BSA. The DNA complexes were visualized by running on a 1x TAE, 1% agarose gel for 1 hour at 70V and then visualized by staining with GelRed (biotium).

### PARP1 purification

Recombinant PARP1 carrying an N-terminal hexahistidine AviTag was produced using a pFastBac vector based Baculo virus expression system for expression in Sf21 insect cells. The cells were harvested by centrifugation and pellets were solubilized in binding buffer (25 mM Tris-HCl, pH 7.4, 150 mM NaCl, 1 mM TCEP, 10 mM imidazole) prior to sonication in the presence of DNase I (Sigma D4527) and EDTA-free protease inhibitor cocktail (Roche 37378900). The lysed pellet was clarified by centrifugation, the supernatant was mixed with 3 mL of NiNTA resin (Thermo Scientific 88222) and incubated for 1 hour at 4 °C. Ni-NTA beads were loaded onto a gravity flow column and washed with binding buffer. PARP1 elution was achieved by employing an imidazole gradient in binding buffer. Fractions containing PARP1 were further purified by ion exchange chromatography on a HiTrap heparin HP (GE Healthcare) column followed by size exclusion chromatography on a Hi load 16/60 Superdex 200 (Amersham Biosciences) prep grade column. SDS-PAGE gel analysis of the purified PARP1 indicated >90% purity and confirmed PARylation activity in the presence of NAD^+^ and DNA breaks (Figure S12). A dissociation constant K_d_=71nM of PARP1 binding to a DNA dumbbell containing a single-strand break was measured with fluorescence polarization (Figure S13) similar to previous studies^21^.

## DATA AVAILABILITY

The data supporting this manuscript is available from the authors upon reasonable request.

## SUPPLEMENTARY DATA

Supplementary data are available.

## ACKNOWLEDGEMENT

The authors thank Gregory Mashanov for helpful discussions and assistance with microscope software; Lizzie Underwood for assistance with protein expression; Richard Thorogate for technical support; and Alice Pyne for helpful discussions and provision of software for automated AFM analysis.

## AUTHOR CONTRIBUTION

N.A.W.B., P.J.H., T.M.O., M.F., B.W.H. and J.E.M. designed the experiments, K.B. purified the PARP protein, N.A.W.B carried out magnetic tweezers, TIRF experiments and gel analysis, P.J.H. carried out AFM experiments, N.A.W.B., P.J.H., B.W.H. and J.E.M. analysed the data, N.A.W.B., P.J.H., K.B., T.M.O., M.F., B.W.H. and J.E.M. discussed the data and conclusions, N.A.W.B., P.J.H., B.W.H. and J.E.M. wrote the manuscript, all authors commented on and approved the manuscript.

## FUNDING

N.A.W.B. is supported by an AstraZeneca-Crick collaborative grant (FC001632) and also by the Francis Crick Institute, which receives core funding from CRUK (FC001119), MRC (FC001119) and the Wellcome Trust (FC001119). P.J.H. is supported by an EPSRC studentship (EP/ L015277/1). The AFM work was facilitated by EPSRC equipment funding (EP/M028100/1).

## CONFLICT OF INTEREST

The PARP inhibitor olaparib is marketed by AstraZeneca. K.B., T.M.O. and M.F. are employees of AstraZeneca. T.M.O. and M.F. are shareholders of AstraZeneca. N.A.W.B. was funded by an AstraZeneca-Crick collaborative grant (FC001632). The other authors declare no conflict of interest.

