## Supplementary Information for "Single-molecule measurements reveal that PARP1 condenses DNA by loop formation"

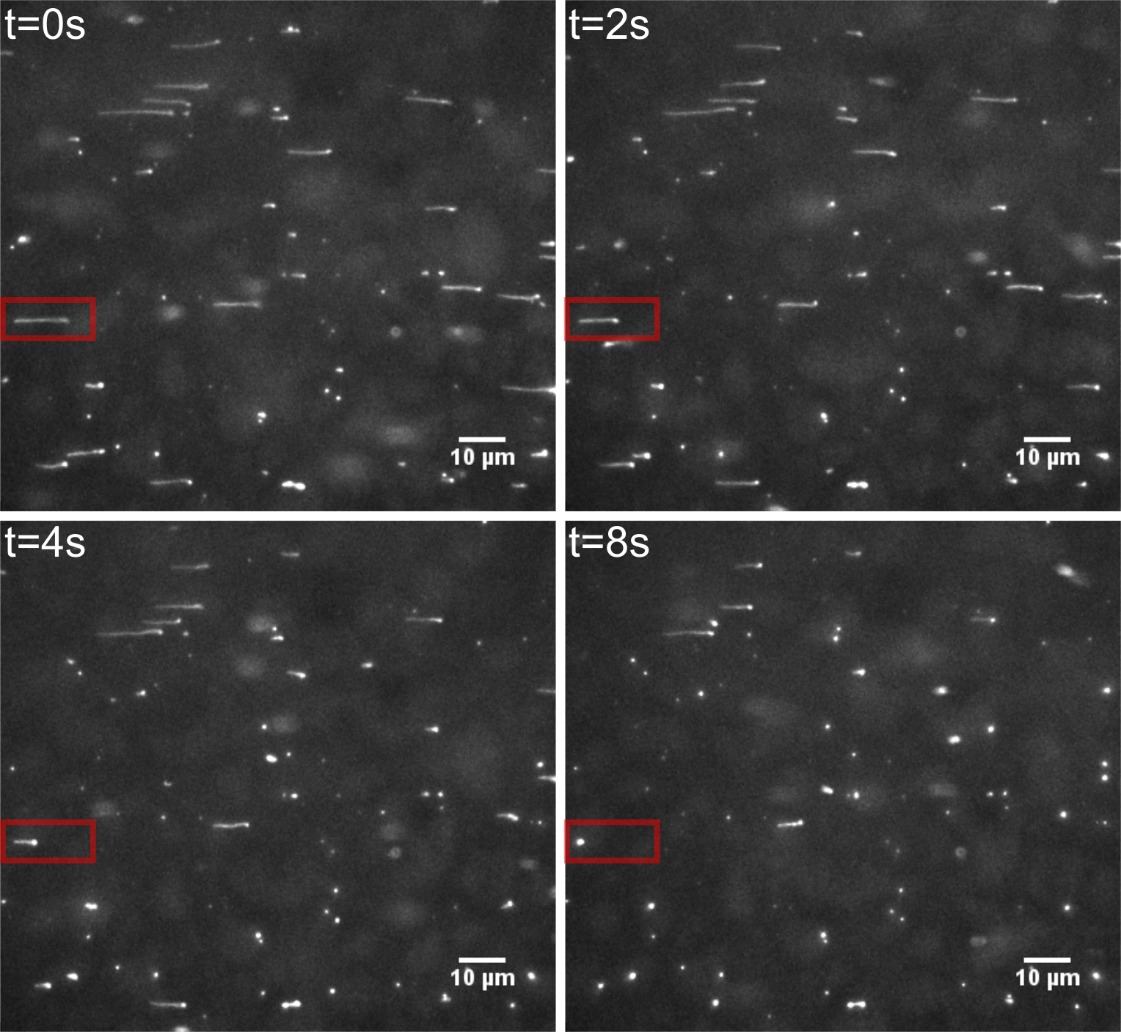

**Supplementary Figure 1.** Full field of view of TIRF image showing DNA condensation after addition of PARP1. PARP1 was estimated to enter the flow cell at t = 0 s based on the dead volume of tubing used for the flow cell and the flow rate of 100 nL/s. Four frames at the indicated timepoints are shown. The red frame highlights the particular DNA molecule shown in Figure 1 in the main text.

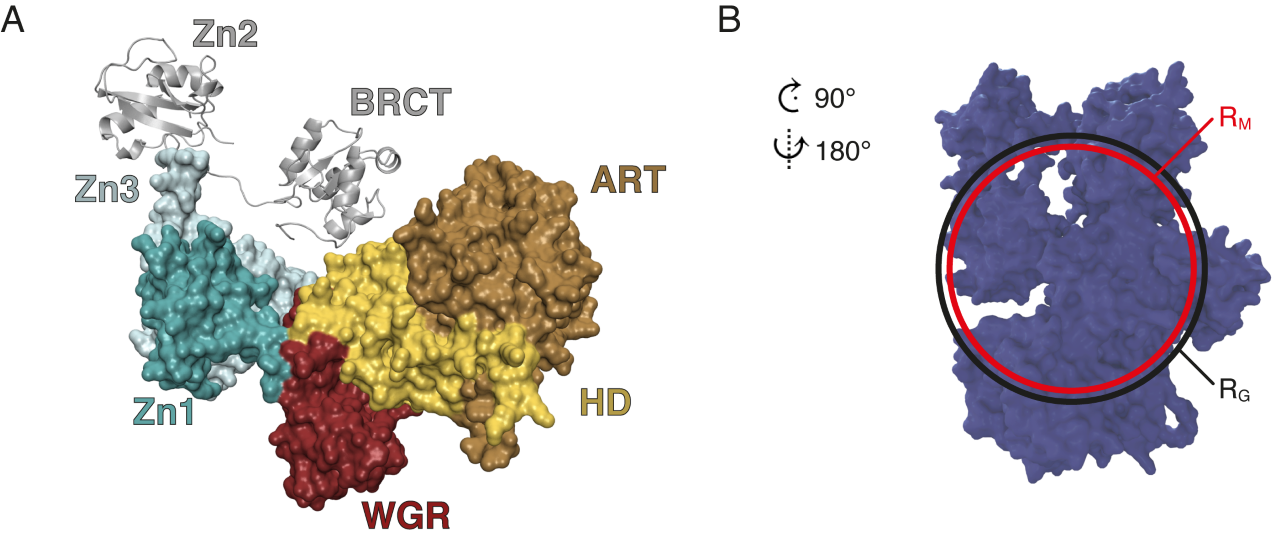

**Supplementary Figure 2.** Estimating PARP1 volume and globularity. (A) Model for the full-length PARP1 crystal structure^1^, generated using the PYMOL Molecular Graphics System (Schrödinger, LLC) from PDB codes: 4DQY, 3ODC(Zn2) and 2COK(BRCT). (B) PARP1 model rotated 90° clock-wise and flipped horizontally with respect to A. The mean radius of crystal coordinates was found to be 3.3 nm with respect to the centre of mass giving a spherical volume of 150 nm^3^. This radius is indicated by the red circle. We also show the radius of gyration (black circle) which has a value of 3.6 nm giving a slightly larger estimate of 195 nm^3^.

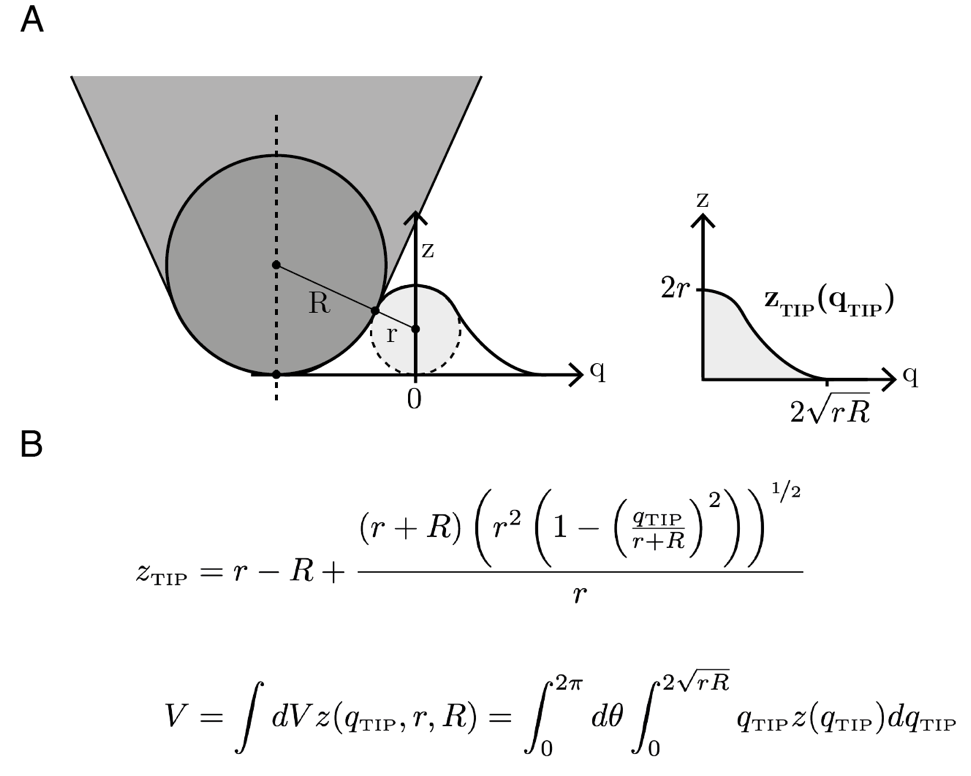

**Supplementary Figure 3.** Geometrical prediction of the volume of proteins as measured by AFM. The protein is assumed to be spherical. (A) Schematic to show how a scanning AFM tip convolutes a molecular profile, *z*_TIP_, where *R* = tip radius, *r* = protein radius and *q*_TIP_ = lateral distance from the tip to the protein (centre to the centre). (B) Expressions for *z*_TIP_, following from geometric arguments outlined previously^2^, and the volume integral expressed in cylindrical coordinates. *R* is estimated to be 3.7 ± 0.8 nm (mean ± s.d.) by considering the measured full-width-half-maximum across four DNA molecules^2^, as obtained by AFM imaging using four different tips, before PARP1 was added. This value of *R* is in close agreement with the nominal tip radius of 5 nm specified by the manufacturer. For these measurements, a setpoint corresponding to a low tip-sample interaction force (<70 pN) was used to minimize compression of the DNA molecule. When probed by the AFM tip, the measured PARP1 radius may be estimated to be *r* = 1.8 ± 0.2 (mean ± s.d, *N* = 11) nm from AFM height data on PARP1 bound to DNA. This indicates the force exerted by the tip has the effect of the PARP1 appearing more compressed, in addition due to possible compaction of PARP1 when binding to DNA. On the other hand, the tip convolution may make the PARP1 appear broader than its actual size (Supplementary Figure 2). Based on this compressed PARP1 radius *r* and the estimated tip radius *R*, a single PARP1 molecule is expected to appear with a volume of *V* = 175 ± 75 nm^3^.

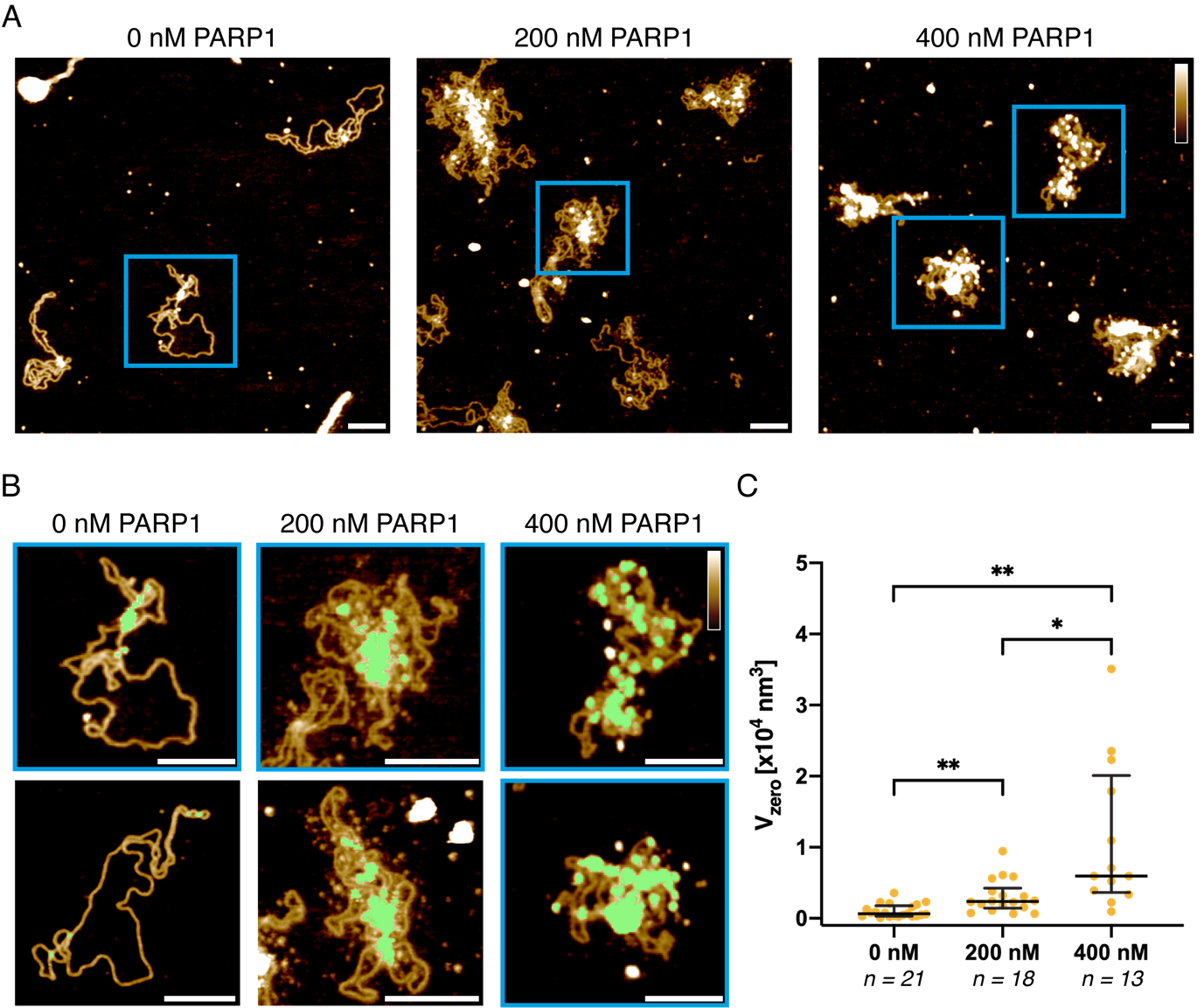

**Supplementary Figure 4.** AFM volume measurements of PARP1 bound to plasmid DNA. (A) AFM images of covalently closed plasmids pre-incubated with 0-400 nM PARP1 *before* deposition on the substrate. Scale bar = 100 nm. Height (colour) scale = 2.5 nm. (B) zoom-in images where the blue squares represent crops taken directly from (A). The green mask indicates the pixels where the local height exceeded a threshold of ~1.5x the height of a DNA molecule, equivalent to ~2 nm above the background. The areas set by the mask were used for volume estimation. Scale bar = 100 nm. Height (colour) scale = 2.5 nm. (C) Boxplots show the observed volume under the masks, V_zero_, measured at different concentrations of PARP1. A shift in the volume indicates the binding of protein to DNA. An unpaired two-sample t-test was performed assuming unequal variances with a significance level of 0.05. **P ≤ 0.01, *P ≤ 0.05, not significant unless stated.

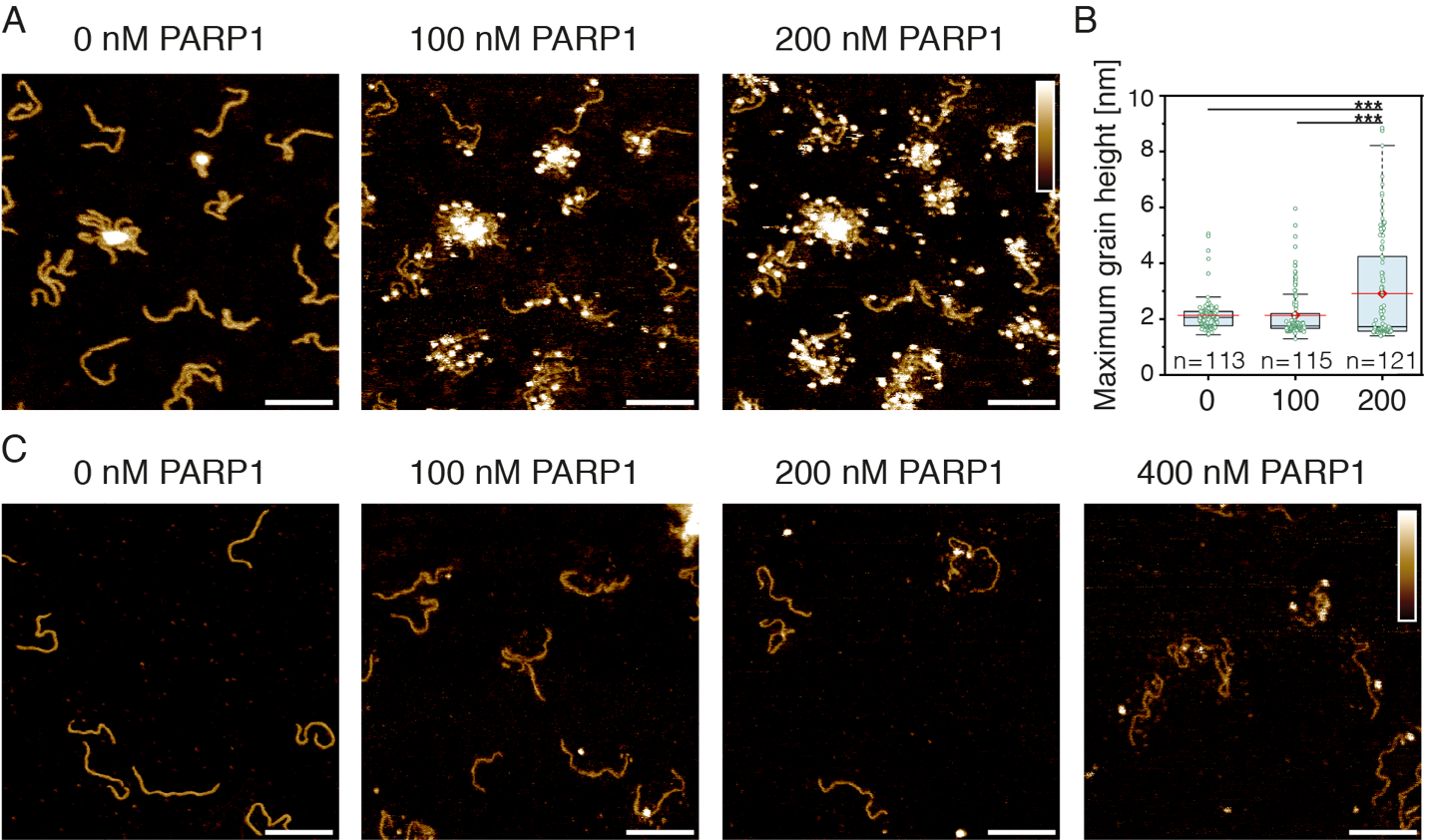

**Supplementary Figure 5.** AFM imaging of PARP1-DNA binding to linear DNA. (A) PARP1 is added to 496 bp linear DNA *after* DNA immobilisation on the AFM substrate and then left to equilibrate for 5 minutes prior to imaging. Images represent the same area with subsequent additions of PARP1 at increasing concentrations. Scale bars: 100 nm; height (colour) scale: 2.5 nm. (B) Boxplots show the maximum grain heights recorded at different concentrations of PARP1, shown in nM, following grain analysis. A shift in the maximum height indicates the binding of protein to DNA. Mean shown in red. An unpaired two-sample t-test was performed assuming unequal variances with a significance level of 0.05. ***P ≤ 0.001, not significant unless stated. (C) PARP1 is pre-incubated with 496 bp linear DNA *before* deposition on the AFM substrate. Scale bars: 100 nm; height (colour) scale: 2.5 nm.

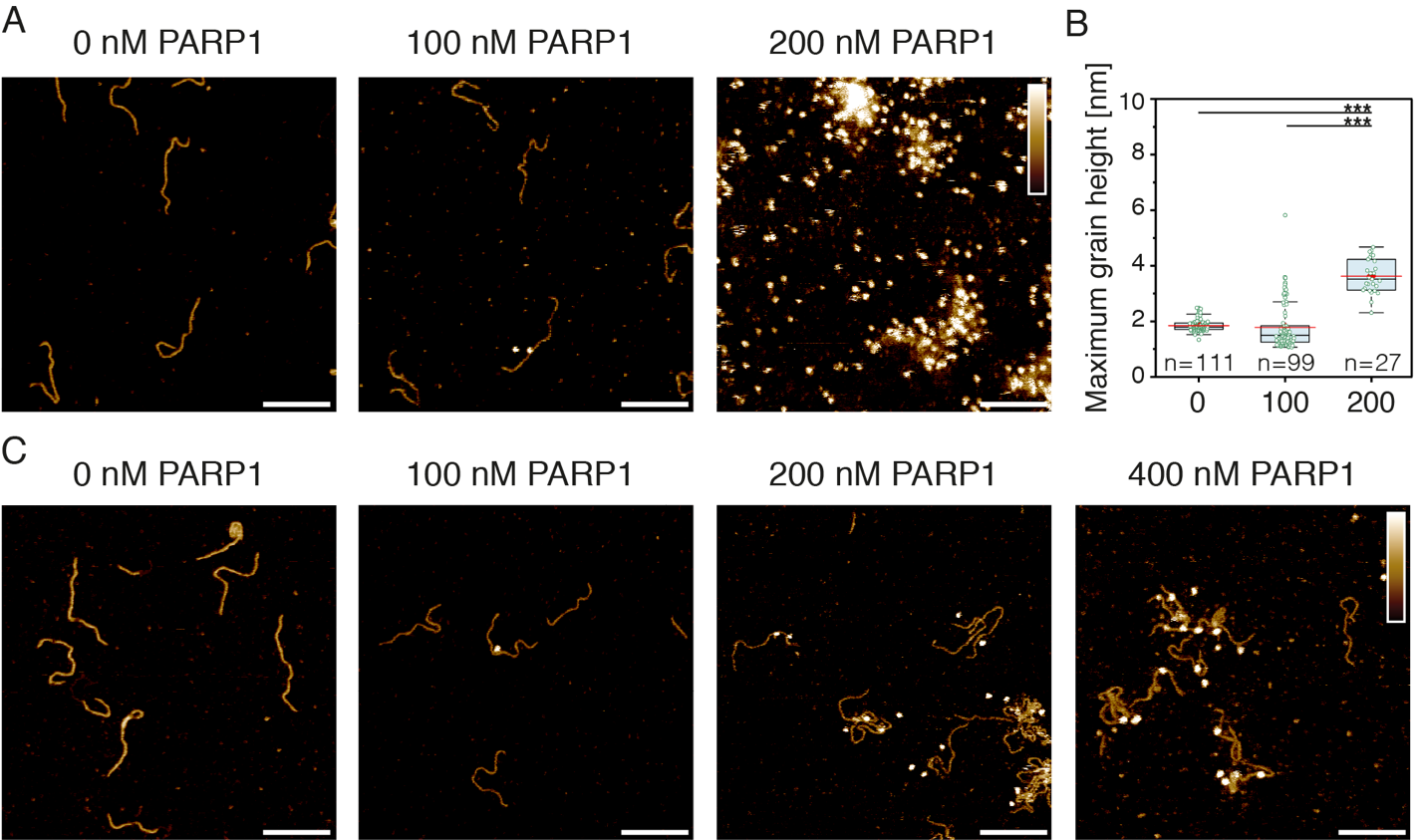

**Supplementary Figure 6.** AFM imaging of PARP1-DNA binding to linear DNA with 500 nM olaparib. (A) 500 nM olaparib was added with PARP1 to 496bp linear DNA *after* DNA immobilisation on the AFM substrate and then left to equilibrate for 5 minutes prior to imaging. Images represent the same area with subsequent additions of PARP1 at increasing concentrations. Scale bars: 100 nm; height (colour) scale: 2.5 nm. (B) Boxplots show the maximum grain heights recorded with 500 nM olaparib at different concentrations of PARP1 (shown in nM). A shift in the maximum height indicates the binding of protein to DNA. Mean shown in red. An unpaired two-sample t-test was performed assuming unequal variances with a significance level of 0.05. ***P ≤ 0.001, not significant unless stated. (C) PARP1 is pre-incubated with 500 nM olaparib and 496 bp linear DNA *before* deposition on the AFM substrate. Scale bars: 100 nm; height (colour) scale: 2.5 nm.

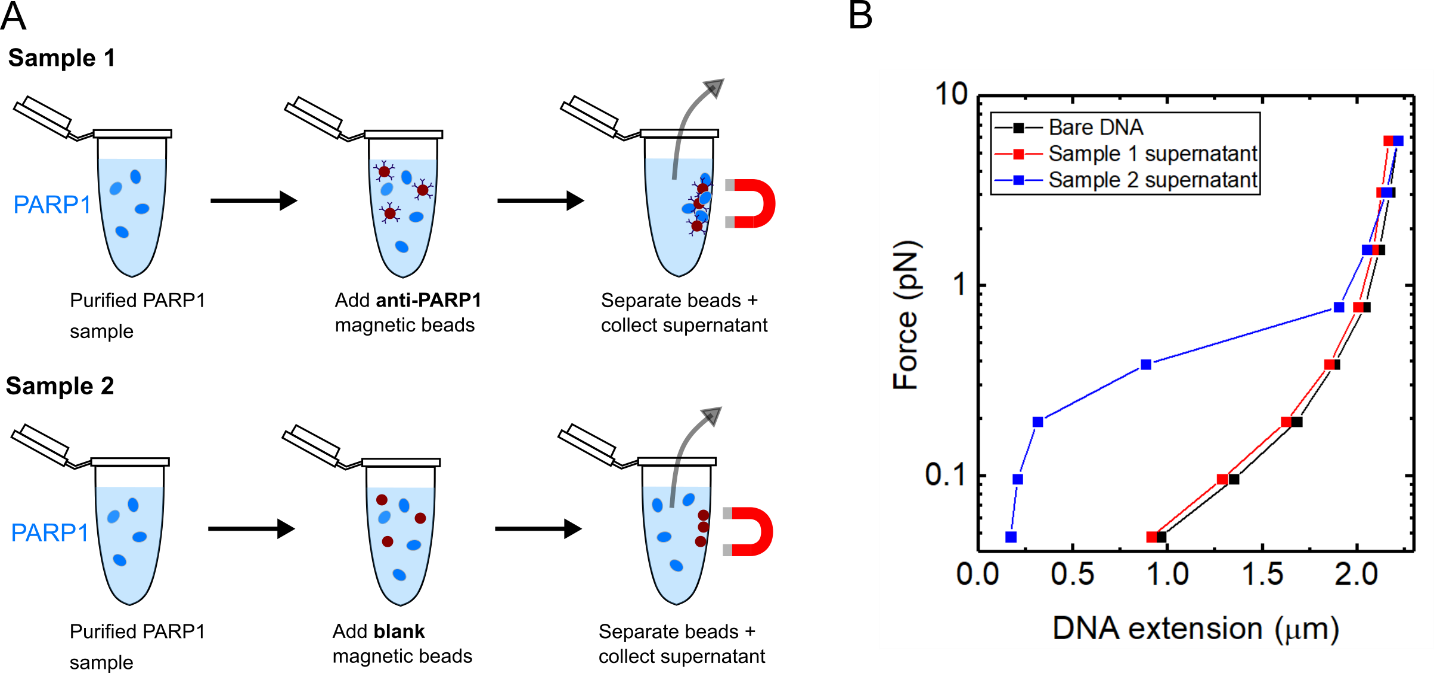

**Supplementary Figure 7.** Immunoprecipitation of PARP1 removes condensation activity. (A) Anti-PARP1 coated magnetic beads were prepared by incubating 8 µl protein G coated magnetic beads (88847, Thermo Fisher) with 8 µl rabbit polyclonal anti-PARP1 (13371-1-AP, proteintech). The beads were washed with buffer A (20 mM HEPES pH 7.8, 150 mM NaCl, 2 mM MgCl2, 0.5 mM TCEP) and resuspended in 8 µl buffer A. For sample 1, 8 µl of 1 µM PARP1 was incubated with 4 µl anti-PARP1 coated beads. For sample 2, the protocol was repeated but the magnetic beads were initially incubated with PBS rather than anti-PARP1 antibodies (this sample acts as a control). For both samples, after incubation for 1 hr with the respective beads, the beads were separated and the supernatant was recovered. The supernatant was diluted 1:1 with 20 mM HEPES pH 7.8, 150 mM NaCl, 2 mM MgCl2, 0.5 mM TCEP, 1 mg/ml BSA, 1 mg/ml β-casein before measurement of force-extension of single 7.9 kbp DNA molecule using magnetic tweezers. (B) Comparison of force-extension for two sample supernatants. The results show that the condensation activity is due to PARP1 rather than potential co-purified contaminants.

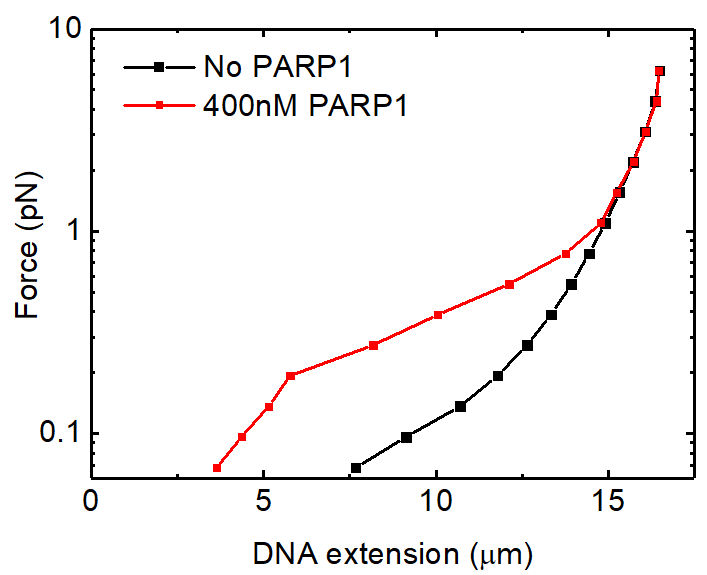

**Supplementary Figure 8.** Magnetic tweezers measurements of full length lambda DNA condensation by PARP1. Lambda DNA was labelled at one end via a biotin labelled oligonucleotide and at the other end by two digoxigenin labelled oligonucleotides. The force was ramped down in steps – each data point shows the average of 10 seconds of trace at each force. Condensation is clearly observed below 1 pN force.

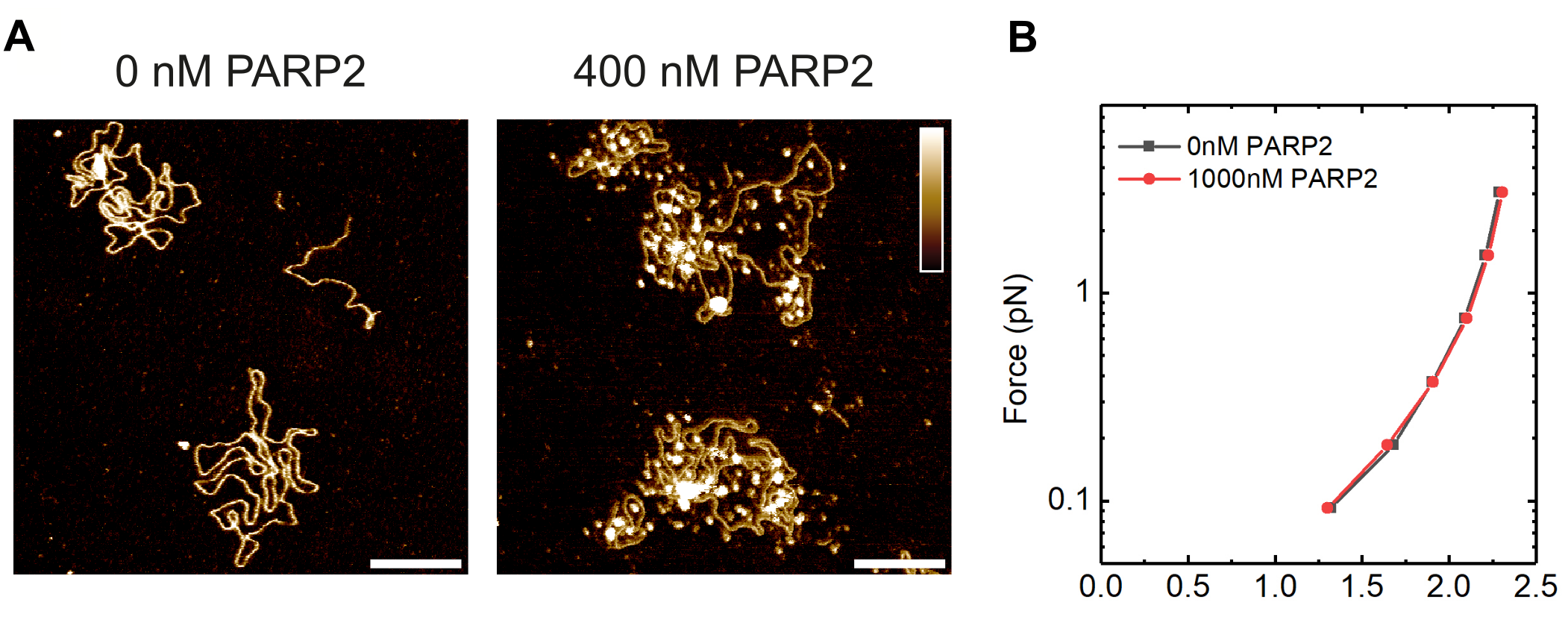

**Supplementary Figure 9.** PARP2 binds to undamaged DNA but does not cause DNA condensation at concentrations up to 1 µM. (A) AFM images comparing plasmid DNA incubated with and without PARP2 (incubation with PARP2 was done before adsorption of the DNA to the AFM substrate). Scale bars: 100 nm; height (colour) scale: 2.5 nm. (B) Force-extension curve for DNA in the presence of 0 nM PARP2 and 1000 nM PARP2. The force was ramped down in steps – each data point shows the average of 10 seconds of trace at each force.

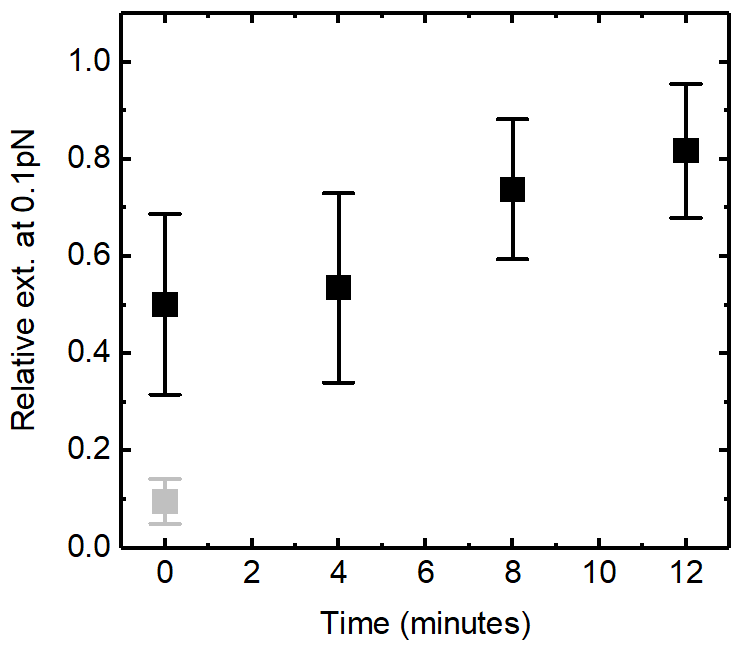

**Supplementary Figure 10.** Loss of condensation for undamaged DNA. The grey data point shows the level of condensation after adding 400 nM PARP1 to undamaged DNA. The timepoints in black show the loss of condensation after flowing through buffer (containing no NAD^+^) and then performing force ramps every four minutes. The data is from N=4 beads (error bars show standard deviation).

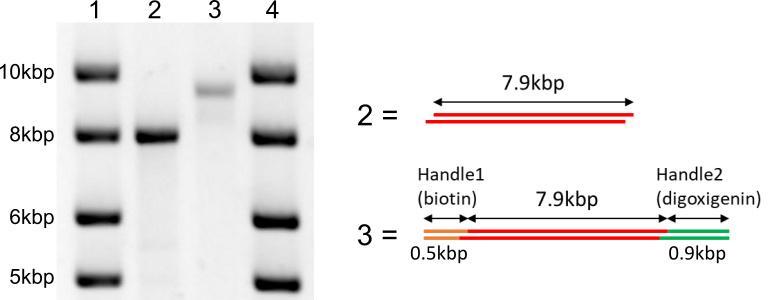

**Supplementary Figure 11**. Agarose gel image of DNA construct used for magnetic tweezers. Lanes 1 and 4 are DNA ladder markers. Lane 2 = Central 7.9 kbp DNA insert purified from lambda DNA. Lane 3 = DNA construct after ligation of two handles.

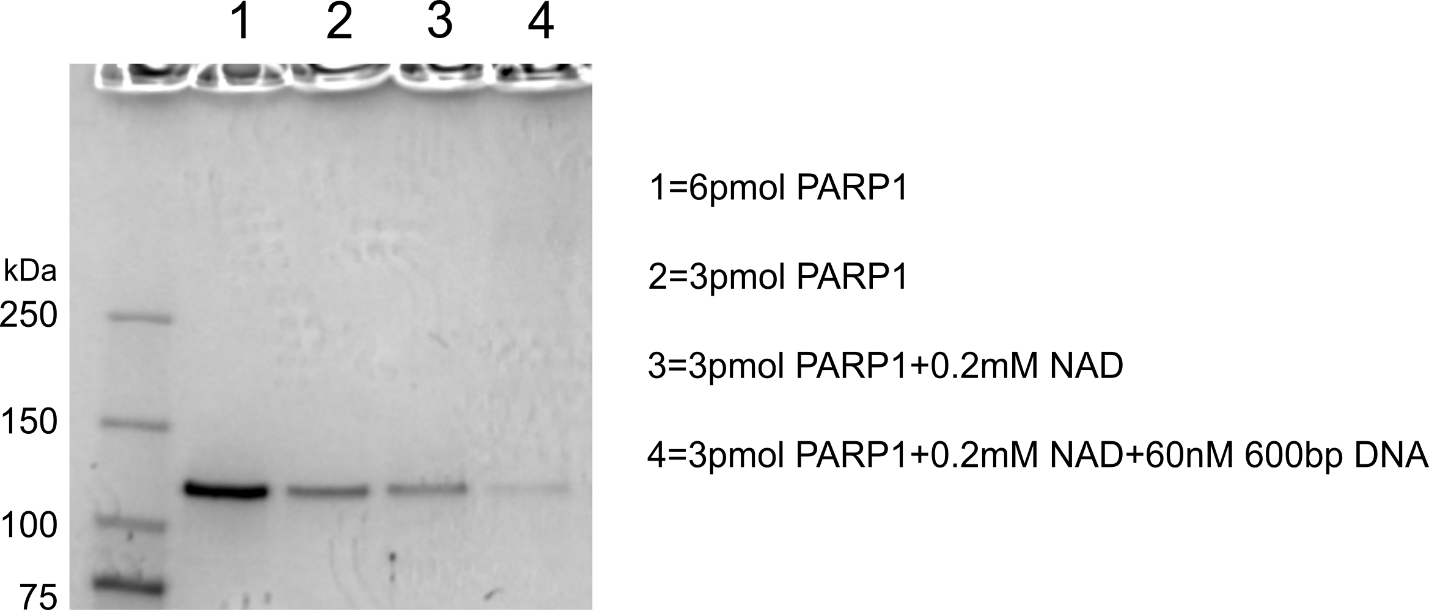

**Supplementary Figure 12.** SDS-PAGE gel analysis of purified PARP1. The samples were incubated for 1 hr at room temperature in a buffer containing 50mM KH_2_PO_4_ pH 7.8, 150 mM KCl, 5 mM beta-mercaptoethanol, 10% glycerol.

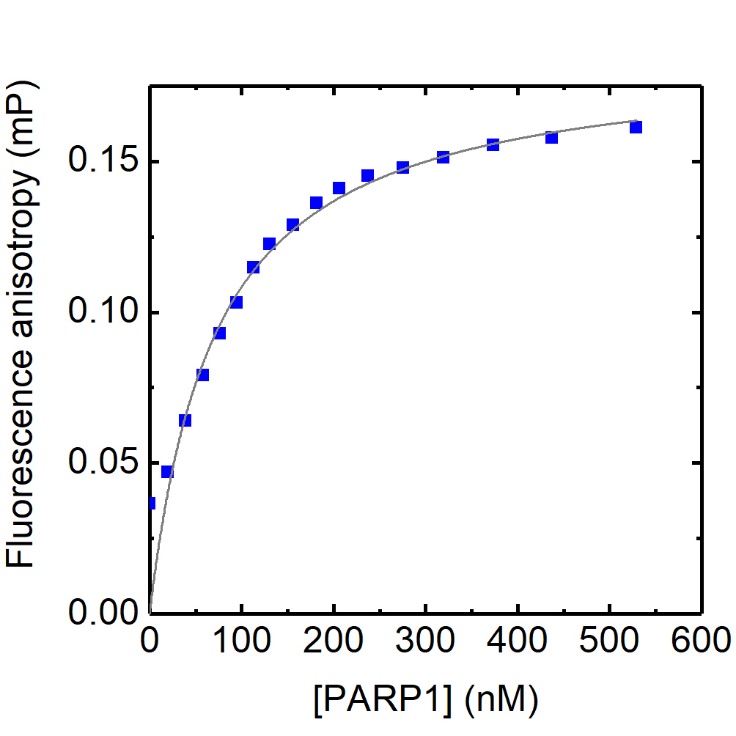

**Supplementary Figure 13.** Fluorescence polarization measurement of PARP1 affinity for a single-stranded break. A DNA dumbbell was formed from the sequence GCT GAG C/iFluorT/T CTG GTG AAG CTC AGC TCG CGG CAG CTG GTG CTG CCG CGA where /iFluorT/ is a fluorescein modification. PARP1 was titrated against an initial concentration of 50 nM DNA. Fluorescence polarization was measured using a Jasco FP-8500 spectrofluorometer with a 100 µL cuvette. The fit is from a one-site binding model to the data. A buffer of 20 mM HEPES pH 7.6, 150 mM NaCl, 1 mM TCEP, 0.005% Brij-35 was used.

1. Langelier, M., Planck, J. L., Roy, S. & Pascal, J. M. Structural Basis for DNA Damage–Dependent Poly(ADP-ribosyl)ation by Human PARP-1. *Science* **728**, 728–733 (2012).

2. Pyne, A., Thompson, R., Leung, C., Roy, D. & Hoogenboom, B. W. Single-Molecule Reconstruction of Oligonucleotide Secondary Structure by Atomic Force Microscopy. *Small* **10**, 3257-3261 (2014).
